# Characterizing the conformational free-energy landscape of RNA stem-loops using single-molecule field-effect transistors

**DOI:** 10.1101/2022.01.13.475405

**Authors:** Sukjin S. Jang, Sarah Dubnik, Jason Hon, Björn Hellenkamp, David G. Lynall, Kenneth L. Shepard, Colin Nuckolls, Ruben L. Gonzalez

## Abstract

We have developed and used high-time-resolution, single-molecule field-effect transistors (smFETs) to characterize the conformational free-energy landscape of RNA stem-loops. Stem-loops are some of the most common RNA structural motifs and serve as building blocks for the formation of more complex RNA structures. Given their prevalence and integral role in RNA folding, the kinetics of stem-loop (un)folding has been extensively characterized using both experimental and computational approaches. Interestingly, these studies have reported vastly disparate timescales of (un)folding, which has been recently interpreted as evidence that (un)folding of even simple stem-loops occurs on a highly rugged conformational energy landscape. Because smFETs do not rely on fluorophore reporters of conformation or on the application of mechanical (un)folding forces, they provide a unique and complementary approach that has allowed us to directly monitor tens of thousands of (un)folding events of individual stem-loops at a 200 μs time resolution. Our results show that under our experimental conditions, stem-loops fold and unfold over a 1-200 ms timescale during which they transition between ensembles of unfolded and folded conformations, the latter of which is composed of at least two sub-populations. The 1-200 ms timescale of (un)folding we observe here indicates that smFETs report on complete (un)folding trajectories in which unfolded conformations of the RNA spend long periods of time wandering the free-energy landscape before sampling one of several misfolded conformations or, alternatively, the natively folded conformation. Our findings demonstrate how the combination of single-molecule sensitivity and high time resolution makes smFETs unique and powerful tools for characterizing the conformational free-energy landscape of RNA and highlight the extremely rugged landscape on which even the simplest RNA structural elements fold.

## INTRODUCTION

In this article, we report a method for tethering a single RNA molecule onto a nanoscopic electrical device that allows us to monitor (un)folding and/or structural rearrangements of the RNA across a broad range of timescales. RNA plays crucial roles in almost all essential biological functions. To perform these functions, RNAs must fold into complex three-dimensional structures and often undergo functionally important conformational changes across a wide range of timescales^1–3^. One of the most fundamental and widely dispersed RNA structural elements are stem-loops, relatively small and fast-folding structures with highly variable thermodynamic stabilities^4–6^ (Figure 1a). Stem-loops commonly serve as folding nuclei for the formation of more complex RNA structural motifs^7^. Additionally, they often participate in vital interactions with other RNAs or proteins and are frequently observed to undergo structural rearrangements that regulate such interactions^1–2, 8–10^. Given such fundamental roles in RNA structure, dynamics, and function, the kinetics of stem-loop (un)folding have long been the subject of extensive characterization using both experimental and computational approaches. Notably, studies using different experimental techniques report disparate stem-loop (un)folding timescales, ranging from 10 μs to 1 s^11–26^, and disagree on whether intermediate states are observed during (un)folding^11, 13–17, 19, 27^. Recent computational studies suggest these inconsistencies originate from the fact that RNAs (un)fold on ‘rugged’ conformational free-energy landscapes containing a large number of energetic wells that are separated by many small and large barriers, and that different techniques simply sample distinct regions of these landscapes^17, 28–30^. As just one example of how this limitation is being addressed by specific experimental techniques, temperature jump (T-jump) experiments are frequently performed at several starting temperatures such that the conformational free-energy landscape can be more extensively sampled^11–14^.

**Figure 1.**
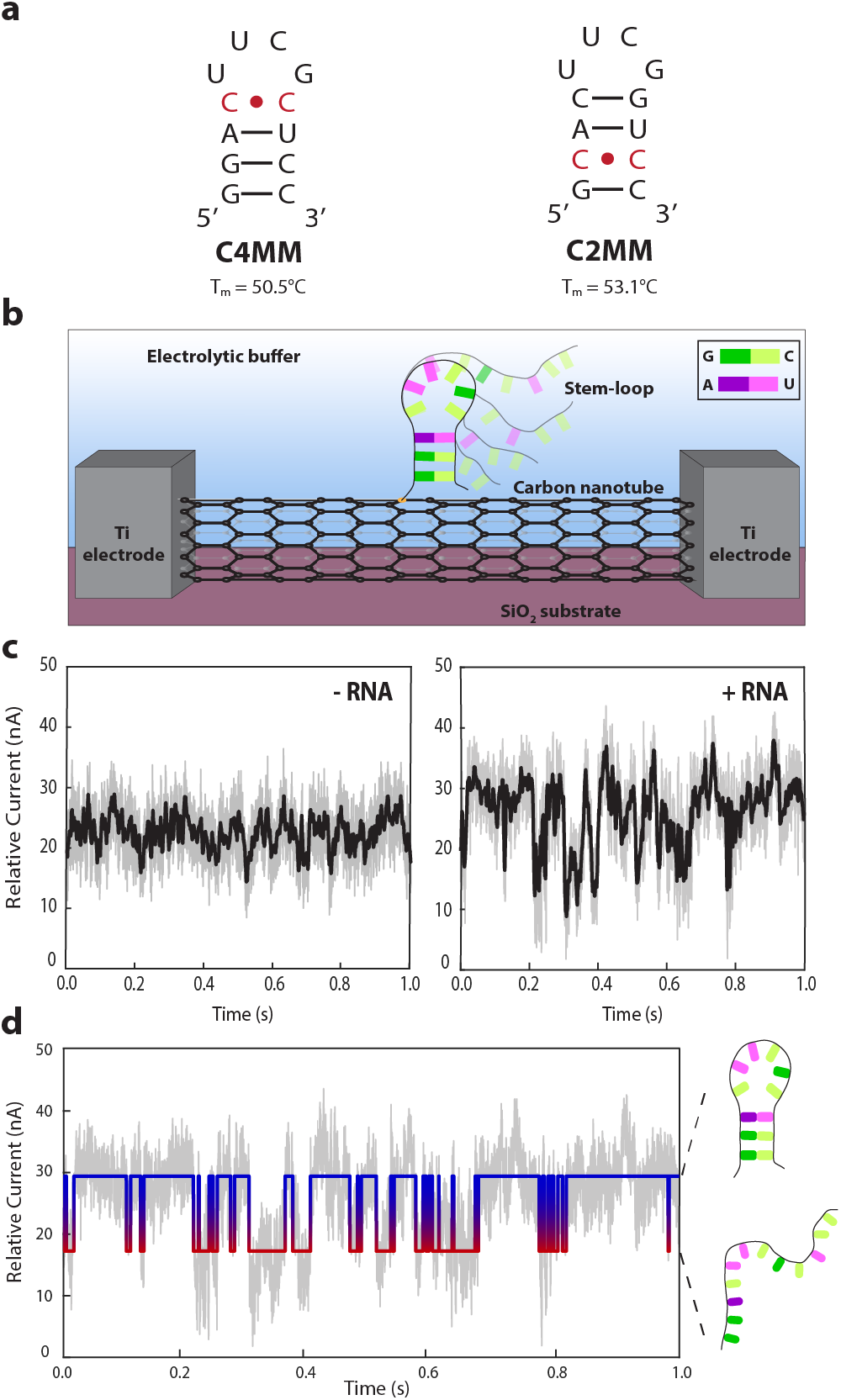
smFET experimental setup. (a) The two stem-loop constructs studied here each introduce a single mismatched bas-pair within the stem of the stem-loop by substituting a cytosine for a guanine. (b) A schematic cartoon of the smFET experimental platform is shown. Each smFET device consists of a SWCNT that serves as a conductive channel between a source- and drain electrode, with a stem-loop construct tethered to the surface of the SWCNT. Details of device fabrication and characterization are described in the Supporting Information and Figure S1. (c) A representative, 1s current versus time trajectory recorded before (left) and after (right) stem-loop tethering. A quantitative analysis of these data demonstrating our ability to distinguish stem-loop-dependent transitions from smFET device noise are provided in Figure S2. (d) An idealized current state versus time trajectory (blue and red), obtained by applying a previously described two-state, drift-corrected, thresholding algorithm^35–36, 42^ is overlaid on the current versus time trajectory. Details of the data analysis procedures are described in the Supporting Information. These trajectories were collected at 200 μs time resolution.

Here, we develop, validate, and demonstrate the use of single-walled carbon nanotube (SWCNT)-based, single-molecule field-effect transistor (smFET) devices for kinetic studies of RNA (un)folding and structural rearrangements. Because current flow through a SWCNT is highly sensitive to the local charge environment, conformational changes of an electrostatically charged biomolecule tethered to the SWCNT surface will manifest as changes in the current flow through the SWCNT^31–34^. Consequently, smFETs enable high-time-resolution, singlemolecule studies of biomolecular dynamics that, uniquely, do not rely on fluorophore reporters of conformational changes nor on the application of mechanical force using an optical trapping instrument or atomic force microscope (AFM)^31–32, 34–41^. Previously, smFETs have been used to make high-time-resolution, single-molecule measurements of DNA hybridization rates^31, 35–37^, G-quadruplex folding kinetics^38^, and protein conformational dynamics^32,34,39–40^.

Using this approach, we report the direct observation of conformational transitions in single RNA molecules across the micro-seconds-to-minutes timescales that are relevant for studies of RNA (un)folding and conformational dynamics. This includes transitions across the 1–200 ms timescales that would be exceedingly difficult, if not impossible, to simultaneously monitor at single-molecule resolution and in a fluorophore- and force-free manner. When the (un)folding of a single RNA molecule is monitored at these timescales and conditions, we find that the thousands of reversible (un)folding events that are observed for even simple stem-loop structures exhibit significant kinetic heterogeneities. Moreover, our results strongly suggest that at least one source of these heterogeneities is the ability of the RNA to arrive at either the natively folded conformation or several misfolded conformations as it folds along a rugged conformational free-energy landscape. Collectively, the results of our studies demonstrate that smFET is a unique and powerful tool for characterizing the conformational free-energy landscapes that govern RNA (un)folding and structural rearrangements.

## RESULTS AND DISCUSSION

### smFETs can monitor the (un)folding of a single RNA stem-loop across a microseconds-to-minutes range of timescales

We designed two RNA stem-loop constructs as the targets of our study, each composed of a four base-pair stem and a well-characterized, thermodynamically stable UUCG tetraloop.^43–44^ The stem of each of these constructs contained a single non-Watson-Crick, C•C mismatched base-pair at either the fourth or second base-pair from the 5’ and 3’ ends of the stem-loop (referred to as the C4MM and C2MM constructs, respectively; see Figure 1a with mismatched base-pairs denoted in red). These mismatched base-pairs were engineered into the stems in order to destabilize the constructs such that they would be expected to readily undergo reversible (un)folding reactions within the temperature range accessible to our smFET devices (up to 50 °C). The corresponding melting temperatures (T_m_s) determined from experiments in which the thermal melting of C4MM or C2MM was monitored using ultraviolet (UV) absorption spectroscopy (referred to hereafter as UV melting) are listed beneath each construct. UV melting data are shown in Figure S3 and the thermodynamic parameters obtained from these data are listed in Table S1. The 5’ end of each construct (Table S2) was functionalized with a primary amine group that was used to tether the RNA to the surface of the SWCNTs, as described below and in the Supporting Information.

To tether the constructs to the SWCNT surface of the smFETs, we followed a previously developed and validated protocol to react the 5’-amine-functionalized RNAs with a carboxylic acid succinimidyl ester-derivatized pyrene (pyrene-NHS) that had been pre-tethered onto the SWCNT surface via noncovalent stacking of the pyrene moiety onto the SWCNT surface^32, 34, 39–40, 45–46^ (Figure 1b). The reaction conditions were optimized such that, on average, no more than one stem-loop was tethered to the SWCNT surface of an smFET (discussed in the following section). Tethering of an individual C4MM or C2MM stemloop to a single smFET resulted in single-molecule current versus time trajectories with clear transitions between distinct current states. The presence of distinct current states in stem-loop-tethered devices was apparent when comparing trajectories recorded before and after a stem-loop was tethered to an smFET (Figure 1c, left and right, respectively). Devices treated with pyrene-NHS and containing a tethered stem-loop resulted in trajectories exhibiting temperature-dependent transition frequencies (Figure S4), whereas devices treated with pyrene-NHS but lacking a tethered stem-loop resulted in trajectories that did not exhibit transitions (Figure S5).

To provide a more quantitative assessment, we began by analyzing the trajectories using a two-state, drift-corrected, thresholding algorithm^42^ that has been previously used to analyze smFET data^35–36^. This algorithm separates the observed states into a cluster of high current states (blue portion of the idealized path depicted in Figure 1d) and a cluster of low current states (red portion). Application of this algorithm to three separate 60-s windows from a trajectory recorded at a relatively low temperature of 23-24 °C (T_low_) prior to tethering a stem-loop, one recorded at T_low_ after tethering a stem-loop, and one recorded at a relatively higher temperature of 44-45 °C (T_high_) after tethering a stem-loop allowed the average number of transitions/60 s in each trajectory to be determined. As shown in Figure S2, <26 % of the transitions observed in a smFET containing a tethered stem-loop at T_low_ and <7 % of the transitions observed in a smFET containing a tethered stem-loop at T_high_ are expected to arise from noise that is misidentified as a transition by the algorithm. The relatively higher percentage of transitions that are expected to arise from the misidentification of noise at T_low_ is due to how rarely the stem-loop transitions to the lower current state at T_low_. In other words, while the absolute number of transitions that are expected to arise from the misidentification of noise at T_low_ is the same as that at T_high_, the very low number of total transitions observed at T_low_ results in the higher percentage of transitions that are expected to arise from the misidentification of noise. Collectively, these results demonstrate that >74 % of the transitions at T_low_ and >93% of the transitions at T_high_ observed in the trajectories arise from the tethered stem-loop.

Previous studies utilizing the smFET platform for biomolecular sensing have shown that the naturally occurring, electrostatically charged chemical groups of a single biomolecule tethered to the SWCNT surface of an smFET device locally gate the current transduction through the SWCNT^31–38, 40^. Consequently, as the tethered biomolecule (un)folds or undergoes a structural rearrangement, changes in the positions of the charged groups relative to the SWCNT surface result in changes to the local gating and corresponding conductivity of the smFET device. More specifically, these studies have demonstrated that relatively closer positioning of negatively charged moieties to the surface of the SWCNT results in relatively higher currents^32–36, 40^. Given the negatively charged phosphate groups of our stem-loop constructs, we tentatively assigned the cluster of higher current states that we observed to an ensemble of folded conformations that would position a relatively larger number of phosphate groups closer to the SWCNT surface. Correspondingly, we assigned the cluster of lower current states to an ensemble of unfolded conformations. To confirm these assignments and characterize the (un)folding of the stem-loops, we first sought to validate that the observed trajectories originated from smFETs onto which a single stem-loop construct had been tethered to the SWCNT surface. Subsequently, we used our smFET data to demonstrate that the population of the higher and lower current states for both constructs decrease and increase, respectively, as a function of increasing temperature, as would be expected if our tentative assignments of the higher and lower current states correctly correspond to the folded and unfolded stem-loop conformations. Moreover, calculating thermodynamic parameters based on these tentative state assignments results in parameters that closely match those determined from conventional, ensemble UV melting experiments (see below).

### Trajectories originate from single stem-loops that have been tethered to single smFETs

Several technical challenges have thus far prevented the use of smFET devices for studies of nucleic acid (un)folding and conformational dynamics. Perhaps the most problematic has been the difficulty of ensuring that only a single nucleic acid is tethered to the surface of a single smFET device. Previous efforts have focused on utilizing both non-covalent and covalent tethering approaches to achieve single-molecule tethering^31–32, 34–38, 40, 46^. In the case of non-covalent tethering, while the approach has been successfully demonstrated for tethering of single protein molecules, tethering of single nucleic acid molecules has not been characterized in a quantitative manner^32, 34, 40, 46^. Here, we focus on utilizing a non-covalent tethering approach^34, 46^ and developing a method to identify single nucleic acid molecules tethered to SWCNTs without perturbing the structural and/or electronic properties of the SWCNT.

To characterize the number of nucleic acid molecules tethered to single smFETs and optimize our tethering protocol so as to ensure that only one molecule was tethered to each individual device, we developed a robust, AFM-based method for visualizing smFET-tethered nucleic acid constructs with single-molecule resolution. Evidence of tethering was facilitated by the formation of nucleic acid-protein complexes that functioned as proxies for nucleic acid-only constructs, which are themselves too small to be confidently visualized using contemporary AFM. We then characterized the number, locations, and specificity with which these complexes tethered to an smFET as a function of varying several parameters of our tethering protocol.

We began by validating the specific tethering of pyrene onto SWCNTs using amine-modified gold nanoparticles. These experiments are described in the Supporting Information and Figure S6. Next, we used the pyrene-NHS incubation protocol to tether 5’-amine-modified, single-stranded, tethering DNA (DNA_T_) containing recognition sites for a previously described variant of the EcoRI homodimeric restriction endonuclease (Table S3), in which the glutamic acid at residue position 111 had been mutated to a glutamine (EcoRI^E111Q^).^47^ EcoRI^E111Q^ is incapable of hydrolyzing its target DNA recognition sequence, instead binding to it with an extremely low equilibrium dissociation constant (*K*_d_) of 2.5 fM.^47–48^ Protein purification and characterization are detailed in the Supporting Information and Figure S7. After tethering of DNA_T_ to the smFETs, single-stranded complementary DNA (DNA_C_) was introduced, followed by EcoRI^E111Q^ in the presence of additional DNA_C_. Once hybridized, DNA_T_ and DNAC formed a double-stranded DNA containing three tandem copies of the EcoRI^E111Q^ recognition site, each separated from the other by a 10 base-pair spacer so as to minimize the possibility that binding of one EcoRI^E111Q^ would sterically occlude binding of additional EcoRI^E111Q^s to the remaining sites (Figure 2a). Binding of six copies (three dimers) of the 31-kDa EcoRI^E111Q^ thus allowed the triple-EcoRI^E111Q^-dimer-bound, double-stranded DNA to be easily imaged and resolved using AFM (Figure 2b). As expected, control tethering experiments performed in the absence of DNAT, but in the presence of DNAC and EcoRI^E111Q^, did not produce the tethered, triple-EcoRI^E111Q^-dimer-bound DNA complexes that were observed in the tethering experiments performed in the presence of DNA_T_ (compare Figure 2c with Figure 2b).

**Figure 2.**
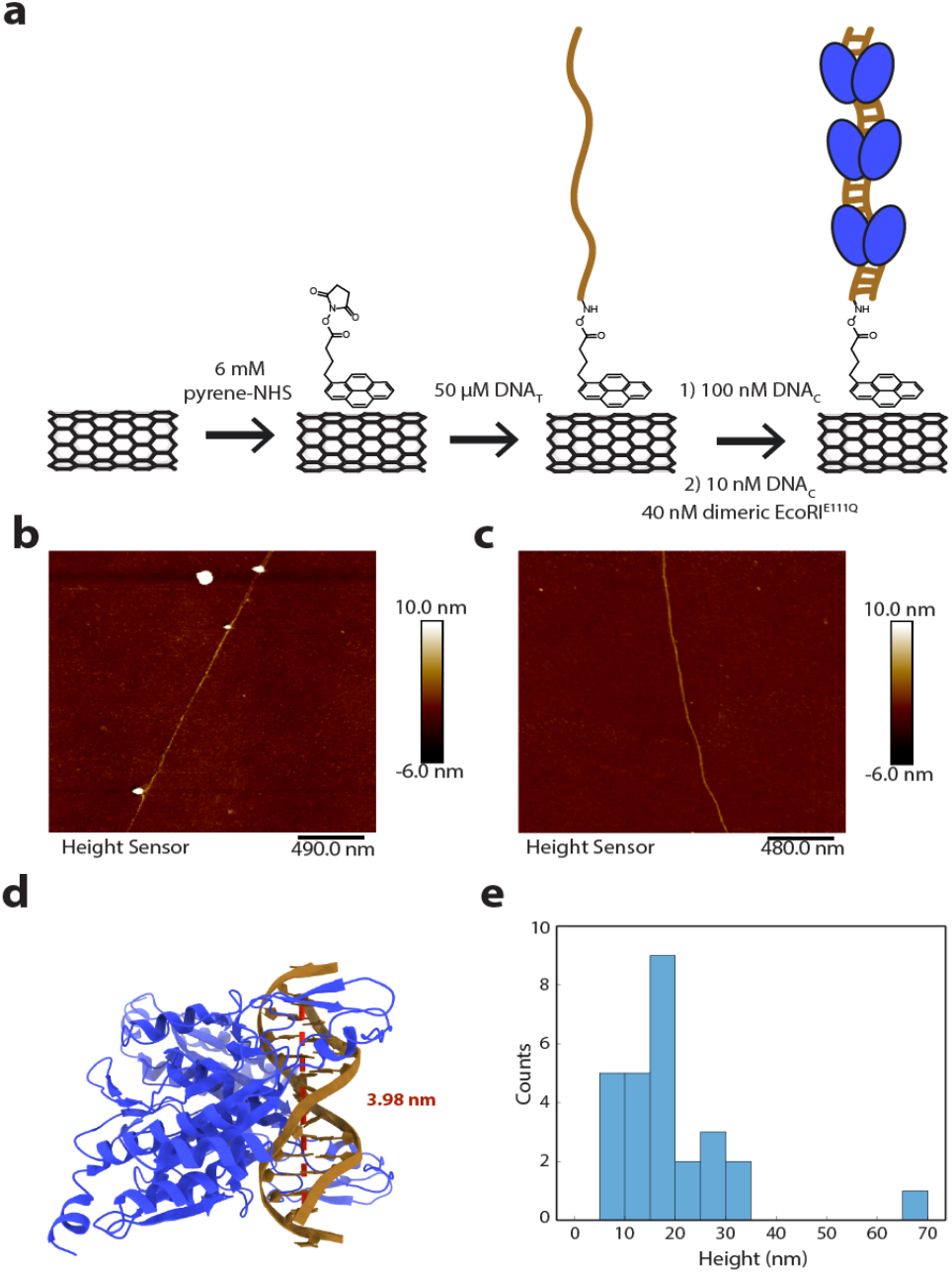
Quantifying and optimizing nucleic acid tethering conditions for smFET experiments. (a) Schematic cartoon of our triple-EcoRI^E111Q^-dimer-bound DNA complex tethering strategy. (b-c) smFET devices were imaged with AFM such that EcoRI^E111Q^-dimer-bound DNA complexes were visible as tall features on individual SWCNTs. (b) Representative AFM image of an smFET device containing 3 tethered EcoRI^E111Q^-dimer-bound DNA complexes generated following the strategy shown in (a). (c) Representative AFM image of an smFET device containing no tethered EcoRI^E111Q^-dimer-bound DNA complexes generated following the strategy shown in (a) with the exception that DNA_T_ was left out of the reaction as a negative control. (d) X-ray crystallographic structure of a single EcoRI^E111Q^-dimer-bound DNA complex (PDB ID: 1ERI), used to determine the length of double-stranded DNA that is occluded by a single EcoRI^E111Q^ dimer. (e) Height distribution of EcoRI^E111Q^-dimer-bound DNA complexes observed in AFM images across three separate experiments.

Given the relatively high concentrations of DNAC and EcoRI^E111Q^ that were used in these experiments (10 nM and 40 nM, respectively), the extremely low *K*_d_ for EcoRI^E111Q^ binding to its recognition sites, and the 10 base-pair separation between each of the three EcoRI^E111Q^ binding sites, we expected to maximize the number of DNATs that would be hybridized with DNA_C_ and harbor three copies of dimeric EcoRI^E111Q^. As shown in the structure in Figure 2d,^49^ a single homodimer of EcoRI^E111Q^ occludes a 3.98-nm, ~12 base-pair length of DNA containing a single recognition site. Considering the persistence length of DNA to be approximately 50 nm,^50^ the 12 base-pair length of DNA was extrapolated to a 48 base-pair, doublestranded B-form DNA helix containing three target sites:

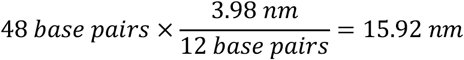

Consistent with this, the length distribution of the individual molecular constructs that were imaged using AFM exhibited a strong correspondence to the expected ~16 nm length of a fully assembled, triple-EcoRI^E111Q^-dimer-bound DNA complex (Figure 2e). We therefore concluded that our AFM-based technique successfully enabled us to quantifiably visualize the number and locations of individual, triple-EcoRI^E111Q^-dimer-bound, DNA complexes, presumably reflecting the number and locations of individual DNA_T_S on the surface of a single smFET device.

Having validated our AFM-based approach for characterizing the number and locations of individual DNA_T_S tethered to a single smFET, we used this method as a readout for varying several of the parameters of our tethering protocol (e.g., the concentration of pyrene-NHS; concentration of 5’-amine-functionalized DNA_T_; incubation times; washing steps and solvents; etc.) to identify the optimal values of these parameters that were most likely to result in tethering of a single DNA_T_ per smFET.

To this end, we found that the optimal conditions included incubating the smFETs in 6 mM pyrene-NHS dissolved in dimethylformamide (DMF), followed by rinsing the smFETs twice with 25 μL of DMF before introducing 50 μM of DNA_T_. These conditions resulted in tethering of approximately one DNA_T_ per 5.79 μm of SWCNT. Given that each smFET is comprised of a SWCNT with a length of approximately 2 μm, a Poisson distribution analysis of tethering under these conditions (see Supporting Information) results in 70.8, 24.4, and 4.2% probabilities of zero, one, and two tethered DNA_T_S, respectively, per device. In other words, given a device with a tethered DNA_T_, there is an 83.9% probability of having a single tethered DNA_T_. Furthermore, the observed distribution of the number of tethering per 2 μm of SWCNT in Figure S8 agrees with the Poisson statistics described here.

### Thermodynamic analyses of the trajectories validate our state assignments

To confirm that transitions between the clusters of high and low current states that we observed report on transitions between ensembles of folded and unfolded stemloop conformations, respectively, we used our smFET devices and optimized tethering strategy to characterize the thermodynamic stabilities, expressed in terms of ΔGs, of the C4MM and C2MM stem-loops. These thermodynamic stabilities were then compared to those determined from conventional, ensemble UV melting experiments to assess the validity of our state assignments.

We began by recording a 10 min long trajectory for each of the C4MM and C2MM constructs at T_low_ as well as T_high_, and analyzing each trajectory with our thresholding algorithm (see Supporting Information). The four trajectories had signal-to-noise ratios (SNRs) of 3-4 and, as expected, all four exhibited transitions between the clusters of high and low current states that were tentatively assigned to ensembles of folded and unfolded conformations, respectively, based on the sensing mechanism of the smFETs. To determine the putative ΔGs of C4MM and C2MM at each temperature, we first quantified the total number of timepoints spent in the presumed ensembles of folded and unfolded conformations (*t*_f_ and *t*_u_, respectively) over a representative, 60-s window of each trajectory and calculated the fractional occupancies of the folded and unfolded conformations (*f*_f_ = *t*_f_ / (*t*_f_ + *t*_u_) and *f*_u_ = *t*_u_ / (*t*_f_ + *t*_u_), respectively) (Figure 3a). These fractional occupancies were then used to calculate the putative equilibrium constants for the folding reactions (*K* = *f*_f_ / *f*_u_) and ΔGs (ΔG = – *RT*lnK, where *R* is the universal gas constant, 1.987 cal/K·mol, and *T* is the temperature at which the measurement was made).

**Figure 3.**
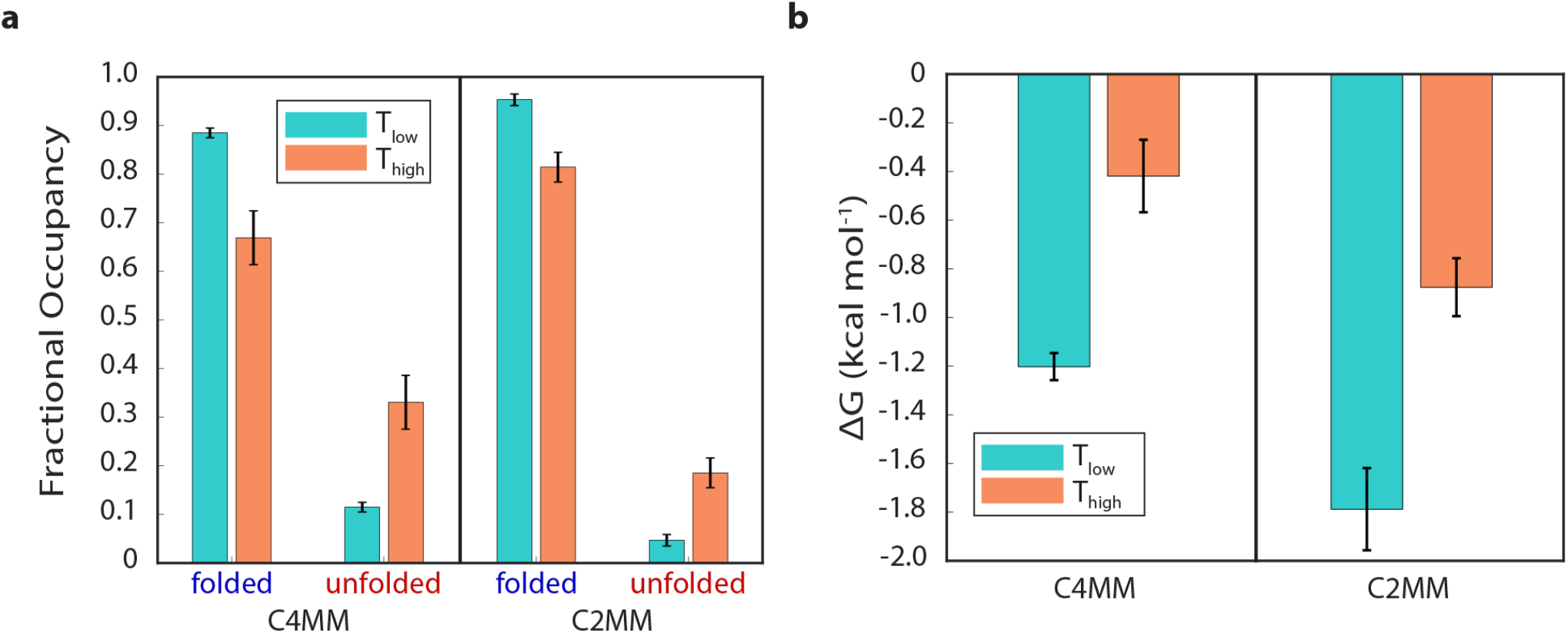
Thermodynamic analyses of C4MM and C2MM current versus time trajectories. (a) Plots of the fractional occupancies of the ensembles of folded and unfolded conformations at T_low_ (23–24 °C) and T_high_ (44–45 °C) for C4MM and C2MM. (b) Plots of the ΔGs of folding at T_low_ (23–24 °C) and T_high_ (44–45 °C) for C4MM and C2MM. Parameters are from the analysis of smFET experiments performed in 100 mM NaCl, 10 mM NaH_2_PO_4_/Na_2_HPO_4_, pH = 7.0 at the specified temperature. Error bars represent the standard deviations of the fractional occupancies and ΔGs calculated from each of three separate 60-s windows of each current versus time trajectory.

As can be seen in Figure 3b and Table S4, the ΔGs for C4MM and C2MM both increase, from −1.20 kcal mol^-1^ and −1.8 kcal mol^-1^, respectively, at T_low_, to −0.4 kcal mol^-1^ and −0.9 kcal mol^-1^, respectively, at T_high_. The fact that the ΔGs for both constructs increase with increasing temperature confirms that we have correctly assigned the clusters of high and low current states observed in the trajectories to ensembles of folded and unfolded stem-loop conformations, respectively; mis-assignment of these states would have resulted in decreases to the ΔGs as a function of increasing temperature, which is not possible.

As expected, and consistent with the results of our smFET experiments, the ΔGs derived from our UV melting experiments on C4MM and C2MM both increase, from −2.48 and −2.7 and kcal mol^−1^, respectively, at T_low_, to −0.59 and −0.74 kcal mol^−1^, respectively, at T_high_ (Table S4). Notably, the ΔGs calculated from our smFET experiments at T_high_ were nearly within error of those calculated from our UV melting experiments (−0.4 ± 0.2 and −0.59 ± 0.06 kcal mol^−1^, respectively, for C4MM and −0.9 ± 0.1 and –0.74 ± 0.05 kcal mol^-1^, respectively, for C2MM; Figure 3b and Tables S4). Similarly, the smFET-derived ΔGs at T_low_ (Table S4) were in good agreement, within ~1 kcal mol^-1^, with the UV melt-derived ΔGs (–1.20 ± 0.06 and –2.48 ± 0.01 kcal mol^-1^, respectively, for C4MM and –1.8 ± 0.2 and –2.7 ± 0.2 kcal mol^-1^, respectively, for C2MM; Figure 3b and Tables S4). Strikingly, the results of the UV melting experiments also corroborate our finding from the smFET experiments that the folded conformation of C2MM is slightly more thermodynamically stable than that of C4MM.

The relatively larger difference between the ΔGs obtained from the smFET and the UV melting experiments at T_low_ is due to the fact that the constructs rarely undergo transitions to the ensemble of unfolded conformations at T_low_. Indeed, the fractional occupancies of the unfolded conformations (f_u_s) of C4MM and C2MM calculated from the UV melting data recorded at T_low_ are f_u_ = 0.02 and f_u_ = 0.01, respectively (Table S4). Due to the small number of excursions to unfolded conformations, even a few instances in which noise within the current states assigned to the folded conformations are misidentified as transitions to unfolded conformations would lead to a significant increase in the ΔGs calculated from the smFET trajectories relative to those calculated from the UV melting data. Indeed, as many as 26 % of the unfolding transitions our thresholding algorithm detects in the trajectories recorded at T_low_ could potentially arise from noise that is misidentified as transitions (see Figure S2).

### Kinetic analyses of the trajectories reveal the ruggedness of the stem-loop (un)folding free-energy landscape

A powerful advantage that smFETs bring to studies of RNA dynamics is that they enable interrogation of the conformational trajectories of individual RNA molecules at an unprecedentedly high time resolution without the need for fluorophore reporters or the application of mechanical force. This property of smFETs potentially provides an avenue for detailed characterization of the ruggedness of the RNA conformational free-energy landscape. To probe this landscape for the (un)folding reactions of the C4MM and C2MM constructs, we performed kinetic analyses of our trajectories.

We began by analyzing the kinetics of transitions between the ensembles of folded and unfolded conformations as defined by our thresholding algorithm (see Supporting Information). Specifically, for C4MM and C2MM at each temperature, we determined the rates of folding and unfolding (*k*_f_ and *k*_u_, respectively) by fitting exponential decay functions to plots of the survival probabilities of the ensembles of unfolded and folded conformations, respectively; the decay lifetimes for the ensemble of unfolded and folded conformations (τ_n_ and τ_f_, respectively) were then used to calculate the rates (*k*_f_ = 1/τ_u_ and *k*_u_ = 1/τ_f_). In all four cases (C4MM and C2MM at each temperature), a single-exponential decay was the simplest function that was needed to properly describe each plot of the survival probability of the ensemble of unfolded conformations, yielding folding rates (*k*_f_) in the range of ~150 to ~500 s^-1^ (R^2^ = 0.991–0.947; see Figure 4 and Table 1). The *k*fs for both C4MM and C2MM increased with increasing temperature (from ~200 to ~500 s^-1^ for C4MM and from ~150 to ~200 s^-1^ for C2MM), consistent with the expectation that the rate of biomolecular (un)folding processes should increase with increasing temperature.

**Figure 4.**
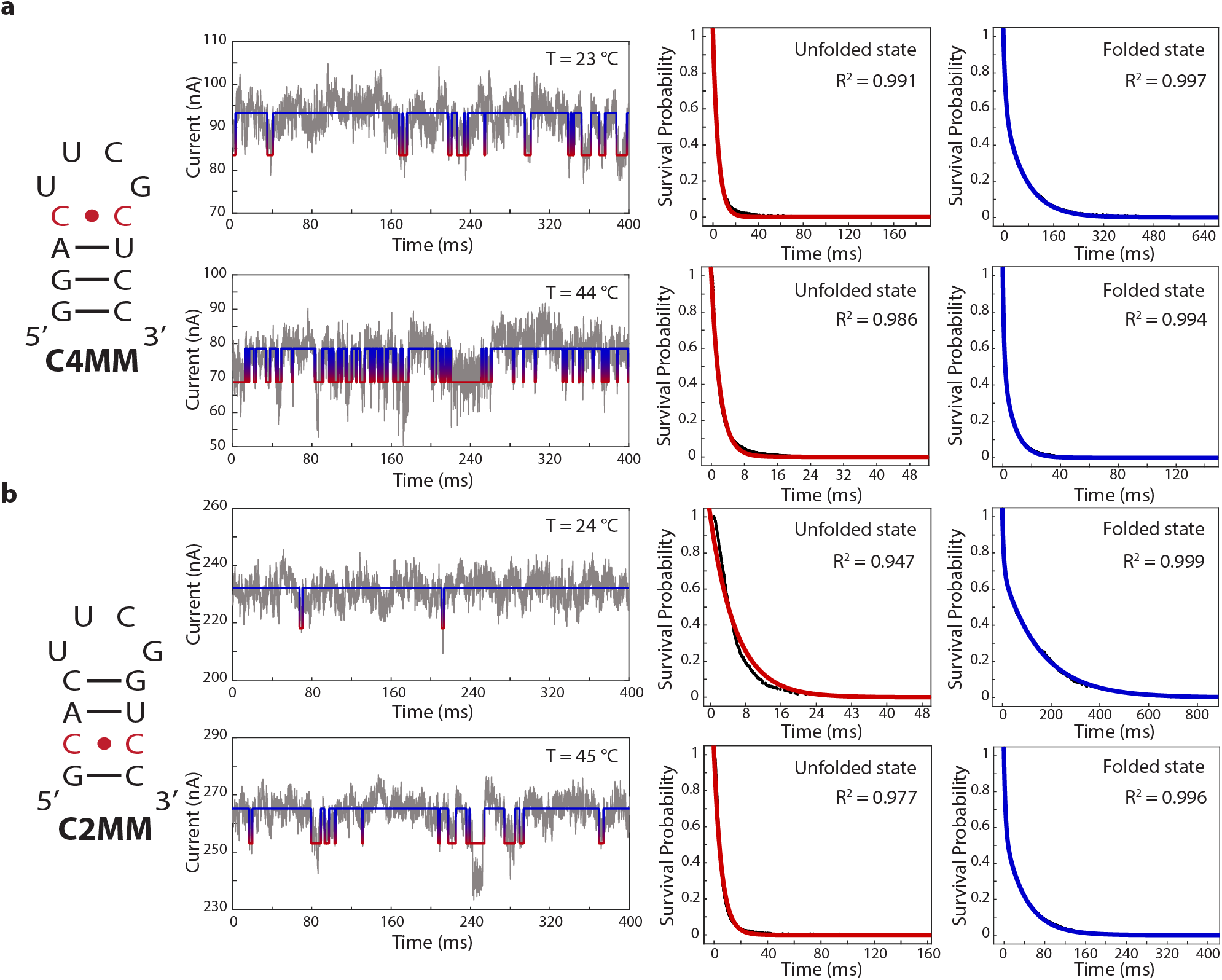
Kinetic analyses of C4MM and C2MM current versus time trajectories. Secondary structure diagrams of the stem-loop constructs with the mismatched base-pairs denoted in red (first column), representative current versus time trajectories (gray) with overlaid idealized current state versus time trajectories (blue and red) (second column), single-exponential fits (red curves) to the survival probability plots of the ensemble of unfolded conformations (black data points) at T_low_ (third column, top) and T_high_ (third column, bottom), and double-exponential fits (blue curves) to the survival probability plots of the ensemble of folded conformations (black data points) at T_low_ (fourth column, top) and T_high_ (fourth column, bottom) for (a) C4MM and (b) C2MM.

**Table 1.** Folding and unfolding rate constants of C4MM and C2MM at the measured temperatures. Rate constants are calculated from the survival probability plots of unfolded and folded state lifetimes.^*a*^

| | Temperature (°C) | $k_f$ (s<sup>-1</sup>) | $k_{u,fast}$ (s<sup>-1</sup>) | Fractional Occupancy (%) | $k_{u,slow}$ (s<sup>-1</sup>) | Fractional Occupancy (%) |
| --- | --- | --- | --- | --- | --- | --- |
| **C4MM** | 23 | 240(29) | 228(78) | 38(3) | 19(5) | 62(3) |
|  | 44 | 486(38) | 983(271) | 45(5) | 146(58) | 55(5) |
| **C2MM** | 24 | 155(23) | 152(26) | 28(4) | 6(2) | 72(4) |
|  | 45 | 160(20) | 315(26) | 41(1) | 22(3) | 59(1) |

<sup>a</sup>Rate constants and (standard deviations) were calculated using the average of three separate 60 s windows from each trajectory.

In contrast, a double-exponential decay was the simplest function that could adequately describe the analogous plots of the survival probability in the ensemble of folded conformations (R^2^ = 0.999–0.994 for double exponential fits vs R^2^ = 0.928– 0.832 for single-exponential fits; see Figure 4 and Figure S9, Supporting Information). Thus, at each temperature, C4MM and C2MM can sample at least two different sub-populations of conformations that our thresholding algorithm collectively assign to the ensemble of folded conformations. In all cases, the longer-lived sub-population that unfolds with the relatively lower unfolding rate (k_u,slow_) in the range of ~10 to ~150 s^-1^ exhibits the higher fractional occupancy, in the range of 55–72% (Figure 4, Table 1). Contrasting with this, the shorter-lived subpopulation that unfolds with the relatively faster unfolding rate (k_u,fast_) in the range of ~150 to ~1000 s^-1^ exhibits the lower fractional occupancy, in the range of 28–45% (Figure 4, Table 1). Again as expected, k_u,slow_ and k_u,fast_ for both C4MM and C2MM increased with increasing temperature (from ~20 to ~150 s^-1^ and from ~10 to ~20 s^-1^ for k_u,slow_ for C4MM and C2MM, respectively, and from ~200 to ~1000 s^−1^ and from ~150 to ~300 s^−1^ for k_u,fast_ for C4MM and C2MM respectively). Moreover, for both C4MM and C2MM, the fractional occupancy of the longer-lived, slower unfolding sub-population decreases with increasing temperature (from 62 to 55% for C4MM and from 72 to 59% for C2MM), suggesting that one of the mechanisms through which ku increases as a function of increasing temperature is via destabilization of the most stable sub-population of folded conformations.

A possible interpretation of the identities of these sub-populations of folded conformations can be drawn from previous theoretical studies of RNA stem-loop (un)folding kinetics.^17, 51–53^ Specifically, these studies propose that the rate-limiting step in the stem-loop folding process is the diffusive search across the conformational free-energy landscape for an RNA conformation that is aligned so as to nucleate formation of interactions that are found in the natively folded stem-loop conformation (i.e., native interactions), while also sampling numerous RNA conformations that form non-native interactions. Formation of such non-native interactions can frequently result in misfolded structures that may or may not be able to proceed to the natively folded stem-loop conformation. In the context of our smFET studies, we hypothesize that our shorter-lived, faster unfolding sub-population of folded conformations represents a collection of misfolded conformations in which non-native interactions persist for a relatively short, but detectable timescale before becoming disrupted so as to allow unfolding of the misfolded conformation. Correspondingly, our longer-lived, slower unfolding sub-population of folded conformations represents the natively folded stem-loop conformation. Future smFET studies focusing on comprehensively elucidating the mechanism of stem-loop (un)folding could use variations of the sequence and length of the stem and loop to further confirm the identity of the two sub-populations of folded conformations and characterize how these sub-populations contribute to the mechanism of stem-loop (un)folding.

In order to further integrate our findings into current models of RNA stem-loop (un)folding, we next turned to the set of ensemble T-jump studies of UNCG (where N = any base) stem-loop constructs that were most similar to those used here.^11–14^ To obtain rate constants for conformational transitions from T-jump experiments, the observed relaxation data must be fit to a defined kinetic model. The earliest T-jump experiments in this set of studies were conducted near the T_m_. The relaxation data obtained from these experiments were well described by a single-exponential function. Correspondingly, these data were fit to a two-state model in which the RNA transitions between states assigned as the fully folded and unfolded conformations^12^. In order to more extensively sample the conformational free-energy landscape, however, follow-up T-jump experiments on the same, as well as similar, stem-loops were conducted at starting temperatures ranging from well below to well above the T_m_. While the relaxation data obtained at the T_m_ were still well-described by a single-exponential function, the data obtained below and above the T_m_ minimally required a double-exponential function. Ultimately, the T-jump data collected at all starting temperatures had to be globally fit to at least a four-state model to capture the observed complexity^11, 13–14^. The four-state model that best described the data was a sequential model in which the RNA transitions from the fully folded conformation to the fully unfolded conformation via two short-lived intermediate states^13–14^. Such a multi-state kinetic model is consistent with other experimental and theoretical studies demonstrating that the energy landscape of even the simplest RNA stem-loops are rugged, involving multiple intermediate states that contribute to the (un)folding process^15, 28, 54^.

When interpreting the results of relaxation data obtained from T-jump experiments, it should be noted that the timescale of the kinetic process of interest will depend heavily on the kinetic model that is used to fit the data. For example, fitting the earliest stem-loop T-jump data collected near the T_m_ with a two-state model resulted in a timescale of (un)folding of 1-100 μs^12^. In contrast, fitting the follow-up stem-loop T-jump data collected over a range of temperatures with a four-state model resulted in a timescale of (un)folding of 0.1-100 ms (see Section G and Table S5 of the Supporting Information for a detailed description of how the timescales of (un)folding reactions for the 4-state model were calculated). The timescale of stem-loop (un)folding that we observe using smFET is 1-200 ms, which is in close agreement with that obtained from analyzing the follow-up T-jump data with a four-state model^13–14^. Together with the close correspondence of the thermodynamic stabilities obtained from the smFET- and UV melting experiments, the close correspondence of the (un)folding timescales obtained from the smFET- and T-jump experiments provide strong evidence that our smFET directly reports on the (un)folding transitions of a single stem-loop. Although our trajectories presumably sample the two short-lived intermediate states that were observed in the follow-up T-jump experiments that were analyzed using a four-state model, the 10-100 μs expected lifetimes of these states (Table S6) and 200 μs time resolution of our smFET experiments would preclude us from directly observing them. Currently, we are developing a complementary smFET platform with improved circuit board design and device architecture that is expected to increase our time resolution by two or more orders-of-magnitude and should allow us to directly detect and characterize such short-lived intermediates in future studies.

Despite having analyzed our data using a simplified, two-state thresholding algorithm, visual inspection of the trajectories reveals that individual excursions into the low- and high-current states that the algorithm collectively assigns as the unfolded and folded conformations, respectively, differ in the absolute value of the observed current. Given the low-frequency ‘1/*f*’ noise of our smFETs^35–36^ and the relatively low SNR of our trajectories, we cannot rule out the possibility that at least some of the variations in the current values within the low- and high-current states are due to these sources of noise. Nonetheless, it is possible that at least some of the excursions to low- and high-current states with different current values represent sampling of distinct unfolded and folded conformations, respectively. As mentioned, ongoing and future improvements of the smFET technology and/or data analysis algorithms (see below) should ultimately allow us to discern additional, authentic conformational states from device noise.

## CONCLUSIONS

We have developed a new, robust, smFET-based approach for single-molecule studies of RNA (un)folding and structural re-arrangements that provides data on timescales of 1–200 ms. Furthermore, we have used this approach to investigate the kinetics of RNA stem-loop (un)folding. Our results provide direct evidence for transitions between ensembles of unfolded and folded conformations and reveal heterogeneities in the unfolding kinetics, suggesting that the folded population consists of natively folded and misfolded conformations. Figure 5 integrates our results and the results of the previous ensemble experimental and computational studies described above as a schematic cartoon of a three-dimensional projection of the multi-dimensional conformational free-energy landscape onto representative RNA stem-loop (un)folding reaction coordinates. As expected from previous studies, the landscape is rugged, featuring many energetic wells corresponding to sets of non-native interactions that act as ‘kinetic traps’ before the RNA samples a nucleating native interaction that allows the RNA to rapidly fold into the stem-loop conformation. In the figure, the dark brown pathway represents a folding event from an smFET experiment in which the RNA initiates from a fully unfolded conformation (Conformation I) and samples a collapsed state containing a nucleating native interaction (Conformation II) that allows it to form the natively folded stem-loop conformation (Conformation III) on a 1–10 ms timescale. We note here that the rate of transitions between Conformations II and III, which can be inferred from relaxation processes observed in ensemble T-jump experiments, are too fast for us to resolve given the timescale of our smFET measurements. The orange pathway represents an unfolding event in which an RNA that has formed non-native interactions and fallen into a kinetic trap (Conformation IV) escapes from the trap on a relatively fast timescale of 1-10 ms to continue its diffusive search across the landscape. The gold pathway, on the other hand, represents an unfolding event in which an RNA that has formed the natively folded stem-loop conformation unfolds on a relatively slow timescale of 10-200 ms.

**Figure 5.**
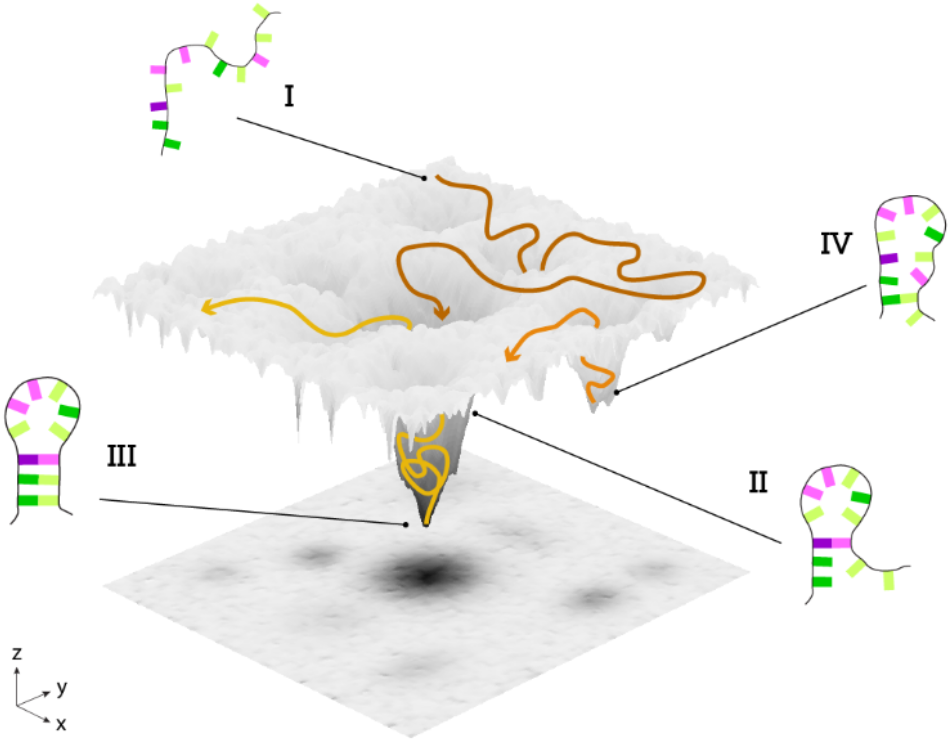
Schematic cartoon of a three-dimensional projection of the multi-dimensional conformational free-energy landscape onto representative RNA stem-loop (un)folding reaction coordinates. Arrows on the landscape depict three representative pathways of (un)folding. The X- and Y dimensions represent reaction coordinates representing different stem-loop conformations and the Z dimension represents the conformational free energy. See Supporting Information for details.

When taken together, the results of our experiments and those performed in the previous experimental and computational investigations highlight the complementary nature of results obtained from these different techniques. The sensing mechanism and single-molecule nature of smFET experiments, for example, uniquely enable direct observation of individual, complete folding trajectories that start from fully unfolded conformations and end in the natively folded conformation, trajectories that exhibit relatively slow kinetics. They also uniquely allow detection and parsing of molecular sub-populations that exhibit different folding pathways. The high time resolution of ensemble T-jump experiments, on the other hand, uniquely enable inference of short-lived intermediate conformations and the rate constants that connect them to the unfolded and folded conformations. Ongoing and future development of the smFET technology described here should enable even more powerful studies of RNA (un)folding and conformational dynamics. An important direction, for example, is the development of smFETs with improved circuit board design and device architecture that will push the time resolution to 1–10 μs and enable us to directly observe and characterize short-lived intermediate conformations currently embedded within the observed ensemble of unfolded and folded conformations. Such developments also promise improvements in the 1/*f* noise and SNRs of smFETs, which will enable the development of more sophisticated, probabilistic data analysis methods that will enable more comprehensive analysis of the resulting current versus time trajectories. Similarly, ongoing and future extensions of the proof-of-principle RNA stem-loop (un)folding experiments reported here should allow a mechanistically richer understanding of stemloop (un)folding. Specifically, varying the lengths and sequences of the stem and loop components of our stem-loop should allow us to further characterize the heterogeneity we have identified in the ensemble of folded conformations, elucidating how these variables contribute to the efficiency with which stem-loops fold from their unfolded conformations into the natively folded conformation instead of any of the several misfolded conformations.

Most importantly, the smFET-based experimental framework for single-molecule studies of RNA (un)folding and conformational dynamics we describe here is entirely generalizable. It can be easily extended to investigate the mechanisms of (un)folding of RNA molecules adopting other stem-loop structures, (un)folding of RNA molecules that fold into structures more complex than simple stem-loops, structural rearrangements of RNA molecules that function as conformational switches, and even the interactions of RNA-binding proteins with their RNA targets. In each of these cases, single-molecule studies that simultaneously monitor timescales across microseconds to minutes are expected to provide mechanistic details that have been difficult or impossible to obtain using other singlemolecule or ensemble approaches. Such developments would bring the field of single-molecule biophysics closer to one of its grand challenges: correspondence between the timescales of single-molecule experiments and molecular dynamics (MD) simulations of those experiments. Such correspondence should allow MD-based, atomic-resolution interpretations of the conformational dynamics captured by single-molecule experiments as well as the experimental testing of MD-derived hypotheses of how conformationally dynamic processes contribute to RNA function.

## ASSOCIATED CONTENT

### Supporting Information

This includes materials and methods used to fabricate and characterize smFET devices, optimize our tethering approach, purify and characterize EcoRI^E111Q^, perform stem-loop UV melting experiments, obtain and analyze stem-loop current versus time trajectories, generate the conformational free-energy landscape figure, and calculate rates of stem-loop (un)folding using previously published T-jump relaxation data are described. Figures illustrate device characterization (Figure S1), quantitative comparison of smFET signals before and after stem-loop tethering (Figure S2), UV melting data (Figure S3), representative current versus time trajectories containing and lacking a tethered stem-loop (Figures S4 and S5), validation of specific tethering of pyrene to SWCNTs (Figure S6), testing of EcoRI^E111Q^ activity (Figure S7), distribution of number of DNA_T_ tethers per device (Figure S8), and additional survival probability exponential curves (Figure S9). Tables report thermodynamic parameters derived from UV melting experiments (Table S1), sequences of RNA constructs used in smFET experiments (Table S2), sequences of DNA constructs used for EcoRI^E111Q^-based tethering controls (Table S3), thermodynamic parameters derived from smFET and UV melting experiments (Table S4), and rate constants calculated using previously published T-jump relaxation data (Table S5,6). This material is available free of charge via the Internet at http://pubs.acs.org.

## AUTHOR INFORMATION

### Author Contributions

All authors have given approval to the final version of the manuscript.

### Notes

K.L.S has a financial interest in Quicksilver Biosciences, Inc., which is commercializing smFET technology for molecular diagnostic applications. Other authors declare no competing financial interest.

## ACKNOWLEDGMENT

C.N. thanks Sheldon and Dorothea Buckler for their generous support. This research was supported by the National Science Foundation (CHE 2004016) and the National Institutes of Health (GM107417). We thank Scott Trocchia for providing the electronic board for the smFETs; Erik Young for providing guidance on building our temperature control system; Kevin Renehan and Eric Pollman for helping with wire-bonding; Riley C. Gentry, Nicholas Ide, and Erik Hartwick for providing guidance during protein purification; Scott Trocchia, Colin Kinz-Thompson, and Korak K. Ray for assistance with data analysis and generation of energy landscape figure. This work was carried out in part in the Clean Room, Electron Microscopy, and Shared Materials Characterization labs of Columbia Nano Initiative (CNI) Shared Lab Facilities at Columbia University, and the Precision Biomolecular Characterization Facility (PBCF). Essential instrumentation in the PBCF was made possible by funding from the U.S. National Institutes of Health (S10OD025102). We thank Jia Ma for management of the PBCF.

---

Authors are required to submit a graphic entry for the Table of Contents (TOC) that, in conjunction with the manuscript title, should give the reader a representative idea of one of the following: A key structure, reaction, equation, concept, or theorem, etc., that is discussed in the manuscript. Consult the journal’s Instructions for Authors for TOC graphic specifications.

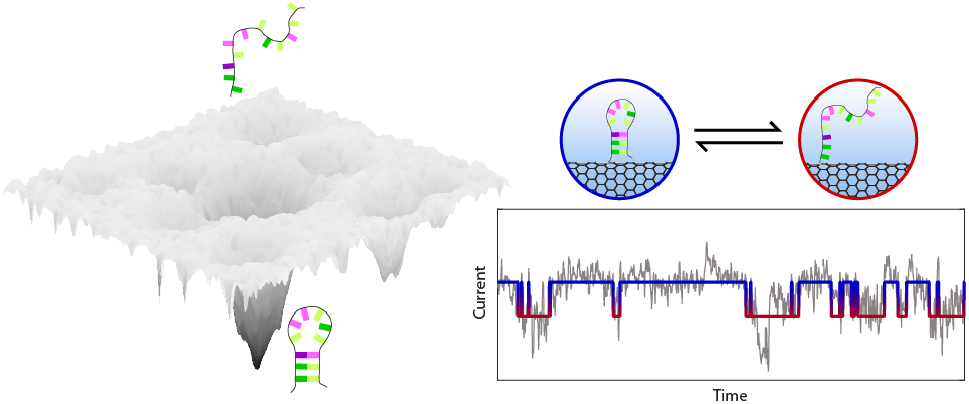

