## Supplementary Information for "Characterizing the conformational free-energy landscape of RNA stem-loops using single-molecule field-effect transistors"

**Table of Contents:**

I. Material and Methods
   A. smFET device fabrication and characterization.
   B. Quantifying and optimizing nucleic acid tethering conditions for smFET experiments.
   C. UV melting experiments on stem-loop constructs.
   D. smFET experiments on stem-loop constructs.
   E. smFET data analysis.
   F. Generating the conformational free-energy landscape figure.
   G. Calculating the rates of stem-loop (un)folding based on a previously described global fit of T-jump relaxation data to a sequential 4-state model.
II. Supplementary Figures
   A. Figure S1: Data from the fabrication and characterization of various, representative smFET devices.
   B. Figure S2: Quantitative comparison of smFET signal before and after stem-loop tethering.
   C. Figure S3: UV melting experiments for C4MM and C2MM.
   D. Figure S4: Temperature dependence of current versus time trajectories recorded using a single, representative smFET device containing a tethered RNA.
   E. Figure S5: Temperature dependence of current versus time trajectories recorded using a single, representative smFET device lacking a tethered RNA.
   F. Figure S6: Validating the specific tethering of pyrene onto SWCNTs.
   G. Figure S7: Characterization of purified EcoRI$^{E111Q}$.
   H. Figure S8: Distribution of the number of tethered DNATs per smFET device.
   I. Figure S9: Single- and double-exponential fitting of survival probability plots of the ensemble of folded conformations.
III. Supplementary Tables
   A. Table S1: Thermodynamic parameters of RNA stem-loop formation.
   B. Table S2: RNA sequences of the stem-loop constructs.
   C. Table S3: DNA sequences used for EcoRI$^{E111Q}$-based DNA tethering controls.
   D. Table S4: Comparison of thermodynamic parameters determined from smFET and UV melting experiments.
   E. Table S5: Overall folding and unfolding rate constants of UNCG tetraloops derived from previous T-jump studies using a sequential four-state model.
   F. Table S6: Individual rate constants of UNCG tetraloops derived from previous T-jump studies using a sequential four-state model.
IV. References

### I. Materials and Methods

All reagents and solvents were purchased from Sigma-Aldrich, Fisher Scientific, or VWR unless otherwise stated. Buffers were prepared with distilled water that was further purified using a Barnstead™ Genpure™ Pro Water Purification System. RNA and DNA oligos (Tables S1 and S2) were purchased from Integrated DNA Technologies (IDT).

#### *A. smFET device fabrication and characterization*

smFET devices were fabricated according to previously described methods.[1-2] Carbon nanotubes (CNTs) were grown *via* chemical vapor deposition on 1 × 1 $cm^2$ silicon (Si)/silicon dioxide ($SiO_2$) substrates. Si wafers covered with a 285 nm-high layer of $SiO_2$ were purchased from Nova Electronic Materials. Pure anhydrous ethyl alcohol with a purity of ≥99.5% was used as the carbon source and iron particles obtained from commercially purchased horse-spleen ferritin were used as a growth catalyst. CNT growths were verified with a Zeiss Sigma VP Scanning Electron Microscope (SEM). Each Si/$SiO_2$ substrate was patterned with sixty-four pairs of titanium source-drain electrodes, where each electrode was 15 µm wide with a source-drain separation of 2 µm. Patterns were written using Nanobeam nB4 Electron Beam Lithography on a spin-coated bilayer of poly(methyl methacrylate) (PMMA) purchased from MicroChem (molecular weight 495K with 4% anisole, followed by 950K with 6% anisole). Metal deposition was then achieved *via* Angstrom UHV E-beam Deposition. An additional lithography and deposition cycle were used to deposit a platinum pseudo-reference electrode on the top and bottom of each substrate and an additional layer of platinum on the end of each titanium electrode (serving as the bond pad for later wire-bonding). The locations of the CNTs were then mapped with respect to the deposited metal using SEM (Figure S1a) and characterized with AFM (Bruker Dimension Icon AFM) **(**Figure S1b) and Raman microscopy (Renishaw InVia Micro-Raman). One CNT per substrate was selected according to its length, location, and Raman characteristics, prioritizing SWCNTs with no noticeable defects or visible residue under AFM. SWCNTs were confirmed by the characteristics of the tangential mode G-band peaks as well as the number of radial breathing mode (RBM) peaks, and defects were identified by the presence of D-band peaks (Figure S1c-d). The removal of the remaining CNTs, as well as the isolation of the selected SWCNT into sixty-four individual devices, was achieved using oxygen ($O_2$) plasma reactive-ion etching (RIE, Plasma Etch) of CNTs exposed *via* electron beam lithography patterning on a single layer of PMMA. Isolation was also confirmed by SEM. The isolated devices were then screened by measuring the current-voltage (I-V) characteristics in air using a semiconductor parameter analyzer (Agilent 4155C). For each device, the I-V curve was obtained by applying a constant source-drain voltages ($V_{ds}$) and a sweeping back-gate voltage ($V_{bg}$) from –10 V to +10 V to the Si/$SiO_2$ substrate (Figure S1e).

#### *B. Characterizing and optimizing the number of nucleic acid molecules tethered to an smFET device*

The general framework for the non-covalent tethering protocols were adapted based on previously described methods.[3-5]

*Validating the specific tethering of pyrene onto SWCNTs*

To test the specificity of pyrene tethering to the SWCNT, amine-functionalized, gold nanoparticles purchased from Cytodiagnostics were reacted with pyrene-NHS and tethered to the SWCNT (Figure S3). For the control experiment, the same steps were followed with pyrene-COOH in place of pyrene-NHS. To achieve tethering, the devices were immersed for one hour in a solution of 1 mM pyrene-NHS prepared in methanol. After the incubation, the devices were rinsed with 0.1 %

v/v Tween-20 in methanol to remove excess pyrene in solution and from the chip surface. Immediately following the rinsing steps, the devices were immersed in 200 nM amine-functionalized 10 nm gold nanoparticles in Functionalization Buffer (10 mM sodium hydride ($NaH_2$)/disodium phosphate ($Na_2HPO_4$) and 100 mM sodium chloride (NaCl), pH = 8.4, and 0.025 % v/v Tween-20) for 90 minutes. The devices were then thoroughly rinsed in water and dried for AFM imaging. AFM measurements for these experiments were carried out in scanAsyst mode using a Bruker Dimension Icon AFM and commercially available scanAsyst-air tips (70 KHz resonance frequency, Bruker).

*Purification and characterization of EcoRI$^{E111Q}$*

Constructs encoding the hydrolytically defective EcoRI$^{E111Q}$ were purchased from Addgene (Dr. Paul Modrich, HHMI & Duke University).[6-7] For overexpression and purification, the gene encoding EcoRI$^{E111Q}$ was sub-cloned into the pET28a(+) vector, which introduces a six-histidine (6×His) affinity tag followed by a highly specific tobacco etch virus (TEV) protease cleavage site at the amino (N) terminus of the EcoRI$^{E111Q}$. The plasmid was transformed into BL21 RIPL cells. 4.5 L of a cell culture was grown to an optical density at 600 nm ($OD_{600}$) of ~0.6 in the presence of 50 μg/ml kanamycin and cells were induced by addition of isopropyl ß-D-1-thiogalactopyranoside (IPTG) to a final concentration of 0.5 mM, collected 4 hours after induction, resuspended in 30 mL of Resuspension Buffer (50 mM tris(hydroxymethyl)- aminomethane hydrochloride (Tris-HCl), pH at 25 °C ($pH_{25°C}$) = 7.5; 1 cOmplete™, EDTA-free Protease Inhibitor Cocktail Tablet (Sigma-Aldrich); and 10% sucrose), and frozen in liquid nitrogen. The frozen cells were then lysed using a SPEX SamplePrep 6875D Freezer/Mill Dual Chamber Cryogenic Grinder and the resulting lysate was diluted by the addition of 100 mL of Wash Buffer 1 (20 mM Tris-HCl, $pH_{25°C}$ = 7.5, and 0.5 M NaCl). The lysate was clarified by high-speed centrifugation in a Beckman Avanti J-E Centrifuge using a JA-17 rotor spinning at 16500 rpm for 30 min and subsequently loaded onto a column containing 4 mL of nickel ($Ni^2$)-nitrilotriacetic acid (NTA) agarose resin. The column was washed with 20 column volumes of Wash Buffer 1 supplemented with 20 mM imidazole. Column-bound EcoRI$^{E111Q}$ was subsequently eluted in 15 mL of Washing Buffer 1 that had been supplemented with 300 mM imidazole. Subsequently, the N-terminal 6×His tag was proteolytically cleaved by adding TEV protease to a final concentration of 0.75 μM to the eluted EcoRI$^{E111Q}$ and dialyzing the cleavage reaction against 2 L of Cleavage Buffer (20 mM Tris-HCl, $pH_{25°C}$ = 7.5; 0.3 M NaCl; and 2 mM βME) at 4°C. After overnight dialysis, TEV-protease-cleaved EcoRI$^{E111Q}$ was flowed through the nickel column once more supplemented with 20 mM imidazole. Further purification of the TEV-protease-cleaved EcoRI$^{E111Q}$ lacking the 6×His tag was achieved by loading it onto a HiTrap Q HP anion exchange column (Cytiva) and eluting it using Anion Exchange Buffer (20 mM Tris-HCl, $pH_{25°C}$ = 7.5, and 0.15 M NaCl) at 4°C. The resulting 15 mL of anion exchange-purified EcoRI$^{E111Q}$ was dialyzed against 2 L of Storage Buffer (40 mM Tris-HCl, $pH_{25°C}$ = 7.5; 300 mM NaCl; 10 mM βME; 0.1 mM ethylenediaminetetraacetic acid (EDTA); 0.15% Triton-X; and 15% glycerol) overnight at 4°C. The resulting 20 mL of dialyzed EcoRI$^{E111Q}$ was then pipetted into 500 μL aliquots and stored at –80 °C until further use. The monomeric concentration of the final, purified EcoRI$^{E111Q}$ was 1.8 μM, as quantified by generating a bovine serum albumin (BSA) concentration standard curve on a sodium dodecyl sulfate-polyacrylamide gel electrophoresis (SDS-PAGE) and comparing the total intensity of the EcoRI$^{E111Q}$ gel band to the total intensities of the BSA standards using ImageJ software. Finally, the ability of our purified EcoRI$^{E111Q}$ to actively bind to its recognition site on double-stranded DNA was characterized using a previously described EcoRI digestion protection assay.[8-9] The results of this assay demonstrate that binding of our purified EcoRI$^{E111Q}$ to λ-DNA protects the 5 EcoRI recognition sites located in the λ-DNA from being recognized, bound, and nucleolytically cleaved into 6 λ-DNA fragments of

21.2, 7.4, 5.8, 5.6, 4.9, and 3.5 kilobases (kbs) by wild-type EcoRI (wtEcoRI) purchased from New England Biolabs (NEB).

*Tethering $DNA_T$ to the pyrene-derivatized, SWCNT surface of smFET devices*

Prior to tethering $DNA_T$ to the smFET devices, a solution of 6 mM pyrene-NHS was prepared in *N,N*-dimethyl formamide (DMF) and a 25 µL aliquot of the pyrene-NHS solution was pulled through the flow cell to soak the devices for one hour. After the incubation, the flow cell was rinsed with two separate 25 µL aliquots of DMF, followed by 150 µL of 0.1% v/v Tween-20 in ethanol to remove the excess pyrene-NHS solution from the flow cell. Immediately following the rinsing steps, the devices were soaked in 50 µM $DNA_T$ in Functionalization Buffer for 90 minutes. From 5' to 3', $DNA_T$ contains a primary amine that is capable of forming an amide bond to pyrene-NHS *via* an NHS ester-activated, cross-linking reaction; a six-carbon linker; a five-nucleotide spacer; three consecutive $EcoRI^{E111Q}$ recognition sites, each separated from the other by 10 nucleotides; and a final, five-nucleotide 3'-terminal sequence (Table S2).

The devices were then rinsed with two separate volumes of 150 µL of Wash Buffer 2 (10 mM $NaH_2PO_4/Na_2HPO_4$ and 100 mM NaCl, pH = 7.0, and 0.05% v/v Tween-20). After conjugation of $DNA_T$ to the pyrene-derivatized SWCNT surface, the devices were soaked in 100 nM $DNA_C$ in Recording Buffer (10 mM $NaH_2PO_4/Na_2HPO_4$ and 100 mM NaCl, pH = 7.0) for 30 min to preassemble a DNA duplex. The devices were then incubated in 10 nM $DNA_C$ with 40 nM $EcoRI^{E111Q}$ dimer for 30 minutes in Binding Buffer (10 mM Tris-HCl, $pH_{25°C}$ = 8.0; 150 mM NaCl; and 10 mM $MgCl_2$) to form $EcoRI^{E111Q}$-dimer-bound DNA complexes, with up to three $EcoRI^{E111Q}$ dimers binding to each DNA duplex. Finally, the surface of the SWCNT was characterized using an AFM to monitor the formation of $EcoRI^{E111Q}$-dimer-bound DNA complexes tethered to the SWCNTs. AFM measurements were carried out in the presence of 10 nM $DNA_C$ and 40 nM $EcoRI^{E111Q}$ dimer in Binding Buffer. A Bruker Dimension Fastscan AFM was used in scanAsyst mode with commercially available Fastscan-C tips (300 KHz resonance frequency, Bruker).

*Poisson distribution analysis of $DNA_T$ tethering*

Each of 32 AFM images was collected by scanning a 4 × 4 µm$^2$ area. The majority of the images included either one or zero triple-$EcoRI^{E111Q}$-dimer-bound DNA complex per 2 µm length of CNT, presumably reflecting the tethering of one or zero $DNA_T$s to the pyrene-derivatized, SWCNT surface of an smFET device. To calculate the probability of tethering any of various numbers of $DNA_T$s to an smFET device, the Poisson distribution of $DNA_T$ tethering to the smFET device was defined as

$$P(N = k) = \frac{\lambda^k e^{-\lambda}}{k!}, \qquad \text{Eq. 1}$$

where *N* is the discrete random variable, in this case it is the number of tethered $DNA_T$s per smFET device; *k* is the number of tethered $DNA_T$s per smFET device whose probability of occurring is being calculated; $\lambda$ is the expected value of *N*, given by dividing the length of the smFET device by the observed average distance between tethered $DNA_T$s ($\lambda$ = 2 µm/5.79 µm = 0.345); and *e* is Euler's number, a constant equal to 2.71828.

#### *C. UV melting experiments on stem-loop constructs*

UV melting experiments were carried out using a Chirascan V100 Spectrometer. For each UV melting experiment, a 10 µM sample of the stem-loop was prepared in Recording Buffer. Each sample was pre-heated to 95 °C, followed by gradual cooling to 5 °C before each experiment. UV absorbance was recorded at 280 nm from 5 °C to 95 °C with a ramp rate of 1 °C $min^{-1}$, and data collected every 0.2 °C. Thermodynamic parameters were determined by nonlinear curve fitting a set of parametric equations for a two-state, unimolecular transition[10] using *cftool* in Matlab.

#### *D. smFET experiments on stem-loop constructs*

Our smFET experimental platform, including the details of electrical measurements and temperature controls, has been described in several previous articles.[1-2, 11] For each smFET experiment, the flow cell was initially flushed with Recording Buffer and a current-voltage characteristic (I-V) curve was collected from –0.4 V to +0.3 V along with a real-time recording at a gate voltage of –0.3 V and a source-drain voltage of either 25 mV or 100 mV. After these electrical measurements, the gate bias and source-drain voltage were set to zero and the flow cell was rinsed with water followed by DMF. To tether the C4MM or C2MM stem-loop constructs (Table S1) to the smFET devices under conditions in which an smFET device containing a tethered stem-loop had an ~84 % probability of having a single tethered stem-loop, we used our optimized $DNA_T$ tethering conditions, determined as described in Section I.B., to tether the stem-loops to the smFET devices. RNA oligos for smFET experiments were not exposed to UV light or thermal treatment to prevent covalent modifications/crosslinking of the RNA that can manifest as kinetic heterogeneity[12]. Disparate levels of such heterogeneities across different studies could differentially affect kinetic measurements and lead to discrepancies in reported rates of RNA (un)folding.

After the stem-loop construct to be studied was tethered and the flow cell was rinsed with Wash Buffer 2, the flow cell was further rinsed in Recording Buffer and maintained in this buffer during recordings. Current versus time trajectories were recorded at $T_{low}$ (*i.e.*, room temperature, or 23–24 °C) at a time resolution of 200 μs for 10 minutes at a gate voltage of –0.3 V and a source-drain voltage of either 25 mV or 100 mV. Following the recordings obtained at $T_{low}$ under these conditions, additional 10-minute recordings at $T_{high}$ (44–45 °C), but otherwise under the same conditions, were obtained by turning on the fan inside the heating chamber and adjusting the voltage of the heat capacitor until achieving $T_{high}$ (Figures S2, 3).

#### *E. smFET data analysis*

The low-frequency '1/*f*' noise inherent to smFET devices results in a 'wandering' baseline current[2, 11] that makes it generally difficult to apply a thresholding algorithm or a hidden Markov model to the raw current versus time trajectories[13]. Additionally, the existence of a wandering baseline and large number of transitions between various current states in each of the trajectories makes it difficult to calculate an accurate rolling baseline for the trajectory that might facilitate the use of a thresholding algorithm or hidden Markov model[13]. To overcome these challenges, the trajectories were analyzed using a two-state, drift-corrected, thresholding algorithm that has been previously developed and used to analyze current versus time trajectories obtained using solid-state nanopore devices[13] as well as those obtained using smFET devices[2, 11]. Briefly, this iterative detection algorithm first performs a rolling average over the entire current versus time trajectory by using a 2,000-time-point window to determine a local baseline current value and subsequently rolls the 2,000-time-point window forward in time by 1 timepoint in order to calculate ~1.5 million local baseline current values for the entire trajectory. Because the two stem-loop constructs used here predominantly sampled the higher current state under all conditions studied, the local baseline current values for all of the smFET experiments performed here were generally closer to the higher current state. Thus, in order to assign each timepoint into either the higher- or lower current state, we used the local baseline current value corresponding to each timepoint as a threshold for the higher current state and set a threshold for the lower current state as follows: (*i*) we created a histogram of the standard deviations of the current values within each of the ~1.5 million 2,000-time-point windows; (*ii*) we took the standard deviation at the peak of this histogram as the representative standard deviation of the entire trajectory; (*iii*) we multiplied this representative standard deviation by a factor of 3 or more, depending on the construct and the smFET device used (a factor of at least 3 was used in order to minimize mis-assignment of timepoints due to noise); and (*iv*) we used the product of this multiplication as a negative offset from the local baseline current value at each given timepoint to set a local threshold for the lower current state. Transitions into the lower current state were defined as those in which a timepoint assigned to the higher current state crossed below the local threshold for the lower current state in the next timepoint. Analogously, transitions into the higher current state were defined as those in which a timepoint assigned to the lower current state crossed above the local threshold for the higher current state in the next timepoint. Finally, a new current versus time trajectory was generated by replacing the timepoints assigned to the lower current state with the local baseline current values at those timepoints. This new trajectory is then used to calculate new rolling averages and local baseline current values. Because the current values assigned to the lower current state in the previous trajectory were replaced with the previously calculated local baseline current values in the new trajectory, the new local baseline current values begin to trend toward the current values of the higher current state. These new local baseline current values are then once again used to set the local thresholds for the higher current state and, likewise, negative offsets from these new local baseline current values calculated as described above are used to set the local thresholds for the lower current state. These new local thresholds are then applied to the original trajectory in order to assign each timepoint to either the higher or lower current state. The algorithm then continues to be iterated until the number of transitions converges.

For each raw current versus time trajectory, the current state assignments made as described above were used to generate an idealized current state versus time trajectory. The idealized trajectories were then used to conduct the thermodynamic and kinetic analyses of the stem-loop constructs described in the main text of this article. Briefly, the fractional occupancy of each current state in each idealized trajectory was calculated by quantifying the total number of timepoints spent in each of the current states. The fractional occupancy of each current state was then used to calculate the *K*s and ΔGs as described in the main text of this article. To obtain the $\tau_u$s and $\tau_f$s, dwell times in each of the current states were compiled and used as input for the *ecdf()* MATLAB function to generate a survival probability distribution and the resulting distribution was fit to either a single-exponential decay function of the form

$$P(t) = e^{-\tau t}, \qquad \text{Eq. 2}$$

where *t* is time, or a double exponential decay function of the form

$$P(t) = Ae^{-\tau_1 t} + (1 - A)e^{-\tau_2 t}, \qquad \text{Eq. 3}$$

where *t* is time and *A* is the amplitude, using the built-in *curve fitting tool(cftool)* in MATLAB. The $\tau_u$s and $\tau_f$s were then used to calculate the $k_f$s and $k_u$s as described in the main text of this article.

#### *F. Generating the conformational free-energy landscape figure*

The conformational free-energy landscape shown in Figure 5 was generated using a custom-written Python script, generously provided by Prof. Colin Kinz-Thompson (Rutgers University at

Newark) and Mr. Korak Kumar Ray (Columbia University), and is freely available at https://github.com/GonzalezBiophysicsLab/energy_landscape/blob/main/2022_JACS_smFET.ipynb.

#### G. Calculating the rates of stem-loop (un)folding based on a previously described global fit of T-jump relaxation data to a sequential 4-state model.

In Sarkar *et al.*, the authors globally fit their T-jump data to a sequential 4-state model (Eq. 4) in order to describe the (un)folding of two different tetraloops[14-15]. The four states are simply denoted as 1-4 in this Supplementary Information section. For a detailed description of the model and the structural conformations assigned to states 1-4, please refer to Sarkar *et al.*[14-15].

$$1 \underset{k_{21}}{\overset{k_{12}}{\rightleftharpoons}} 2 \underset{k_{32}}{\overset{k_{23}}{\rightleftharpoons}} 3 \underset{k_{43}}{\overset{k_{34}}{\rightleftharpoons}} 4 \qquad \text{Eq. 4}$$

In their studies, the free energies of the four states ($\Delta G_i$, where *i* denotes a specific state) and the average free energy barriers connecting them ($\Delta G^{\dagger}_{i\leftrightarrow j}$ where *i* and *j* denote neighboring states) were fit to the following linear models:

$$\Delta G_i(T) = \Delta G_i^{(0)} + \Delta G_i^{(1)}(T - T_0), \qquad \text{Eq. 5}$$

$$\Delta G^{\dagger}_{i\leftrightarrow j}(T) = \Delta G^{\dagger(0)}_{i\leftrightarrow j} + \Delta G^{\dagger(1)}_{i\leftrightarrow j}(T - T_0), \qquad \text{Eq. 6}$$

where $T$ is the temperature at which the free energies and average free energy barriers are specified (for the current study, $T$ = 328.15 K), $T_0$ is the reference temperature, $\Delta G^{(0)}$ is the free energy at the reference temperature, and $\Delta G^{(1)}$is the first derivative of the free energy with respective to temperature. The values of $T_0$, $\Delta G^{(0)}$, and $\Delta G^{(1)}$ are provided in Sarkar *et al.*[14-15].

Using the $\Delta G_i$s and $\Delta G^{\dagger}_{i\leftrightarrow j}$s provided by Sarkar *et al.*[14-15], we then calculated the directional free energy barriers using the following relationships:

$$\Delta G^{\dagger}_{ij} = \Delta G^{\dagger}_{i\leftrightarrow j} + \Delta G_{ij}/2 \qquad \text{Eq. 7}$$

$$\Delta G^{\dagger}_{ji} = \Delta G^{\dagger}_{i\leftrightarrow j} - \Delta G_{ij}/2 \qquad \text{Eq. 8}$$

Where $\Delta G_{ij}$ is the free energy difference between states *i* and *j*.

Subsequently, we calculated the rate constant from state *i* to *j* ($k_{ij}$) by plugging each directional free energy barrier at a given temperature into the following equation:

$$k_{ij} = k_m \exp\left[\frac{\Delta G^{\dagger}_{ij}(T)}{RT}\right], \qquad \text{Eq. 9}$$

where $k_m$ is the pre-exponential factor calculated at $T$ using the previously described temperature dependence of $k_m$[14, 16] and R is the universal gas constant (8.314 J $K^{-1}$ $mol^{-1}$).

To determine the rate of unfolding from the fully folded state (State 1) to the fully unfolded state (State 4), we calculated the mean first passage time from State 1 to 4 based on an equation derived in Kinz-Thompson *et al.*[17],

$$1 \underset{k_{21}}{\overset{k_{12}}{\rightleftharpoons}} 2 \underset{k_{32}}{\overset{k_{23}}{\rightleftharpoons}} 3 \overset{k_{34}}{\longrightarrow} 4 \qquad \text{Eq. 10}$$

$$\langle \tau_f \rangle = \frac{1}{k_{12}}\left[1 + \frac{k_{21}}{k_{23}} + \frac{k_{21}k_{32}}{k_{23}k_{34}} + \frac{1}{k_{23}}\left[1 + \frac{k_{32}}{k_{34}}\right] + \frac{1}{k_{34}}\right] \qquad \text{Eq. 11}$$

Similarly, to calculate the rate of folding, the mean first passage time from the fully unfolded state (State 4) to the fully folded state (State 1) was calculated based on the analogous equation derived in Kinz-Thompson *et al.*[17]:

$$4 \underset{k_{34}}{\overset{k_{43}}{\rightleftharpoons}} 3 \underset{k_{23}}{\overset{k_{32}}{\rightleftharpoons}} 2 \overset{k_{21}}{\longrightarrow} 1 \qquad \text{Eq. 12}$$

$$\langle \tau_u \rangle = \frac{1}{k_{43}}\left[1 + \frac{k_{34}}{k_{32}} + \frac{k_{34}k_{23}}{k_{32}k_{21}} + \frac{1}{k_{32}}\left[1 + \frac{k_{23}}{k_{21}}\right] + \frac{1}{k_{21}}\right] \qquad \text{Eq. 13}$$

### II. Supplementary Figures

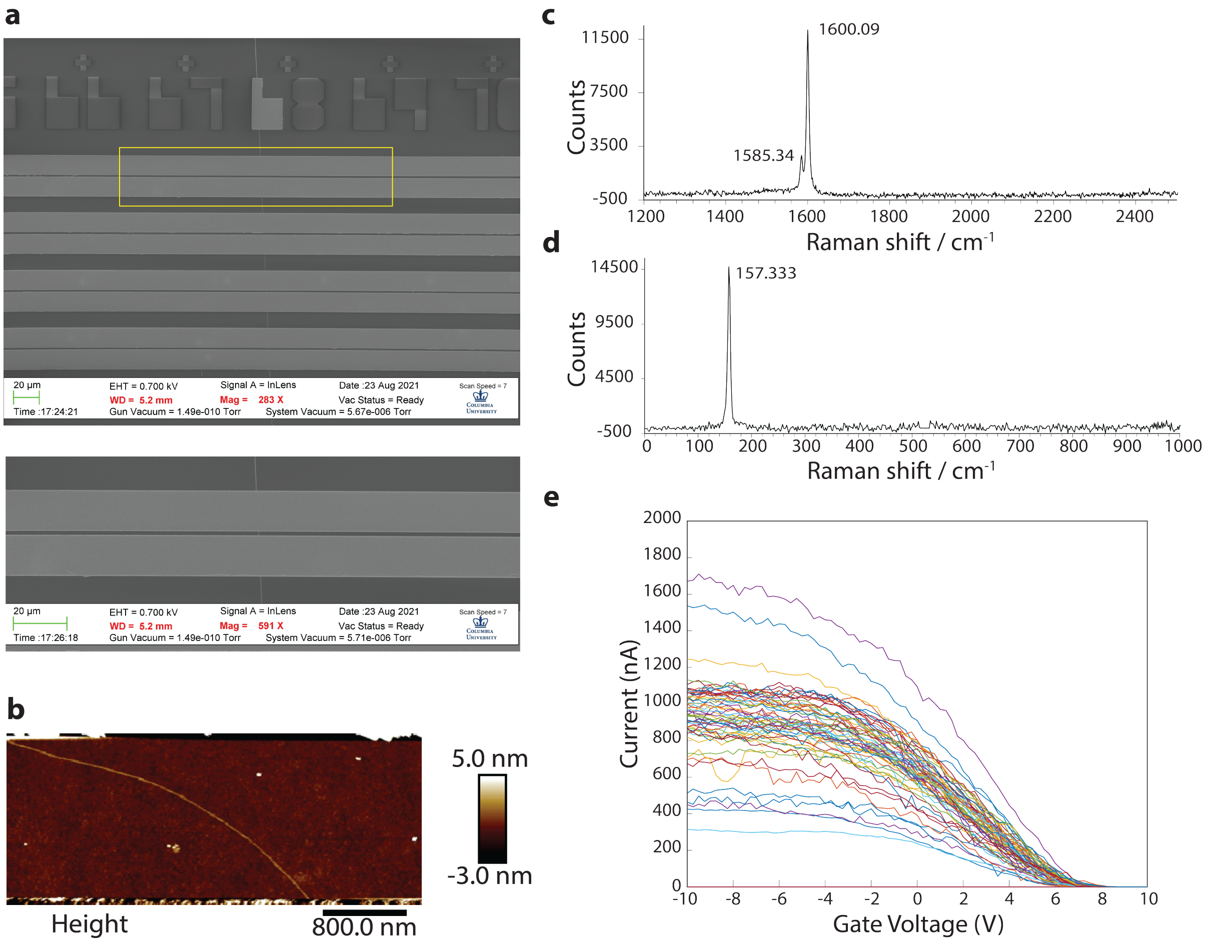


**Figure S1. Data from the fabrication and characterization of various, representative smFET devices. (a)** SEM image of four representative smFET devices, with source and drain electrodes deposited on a SWCNT. One of these smFET devices is shown in a magnified view. **(b)** AFM image of a single, representative smFET device between source and drain electrodes. **(c)** Raman spectrum of a representative CNT showing the G-band peaks. **(d)** Raman spectrum of a representative CNT showing the RBM peak. **(e)** Superimposed I-V curves for all smFET devices on a single $Si/SiO_2$ substrate, after isolation.

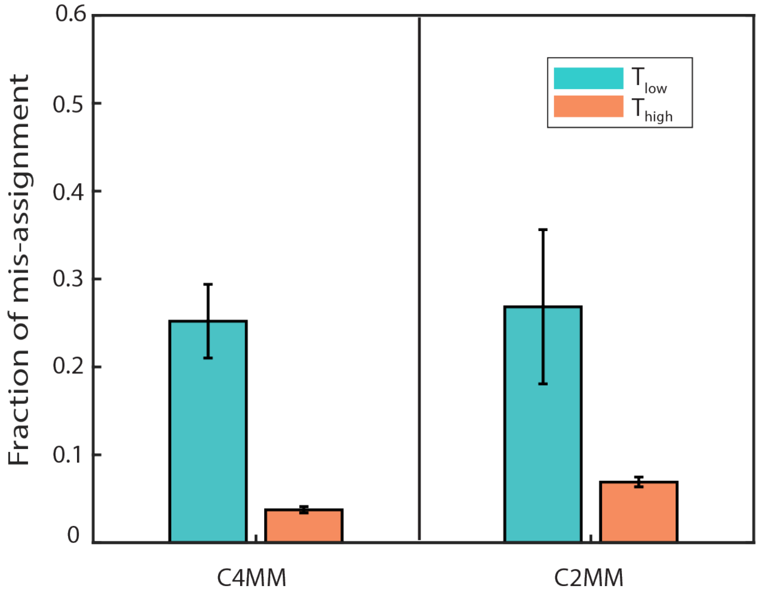


**Figure S2. Quantitative comparison of smFET signal before and after stem-loop tethering.** Fraction of mis-assignments were calculated by comparing the number of transitions observed in a 60s current trajectory prior to tethering a stem-loop at $T_{low}$ (23–24 °C) to the number of transitions observe in a 60s current trajectory after tethering a stem-loop at $T_{low}$ (23–24 °C) or $T_{high}$ (44–45 °C). Error bars represent the standard deviations of three fraction of mis-assignments, each calculated from a pair of 60-s trajectories before and after stem-loop tethering.

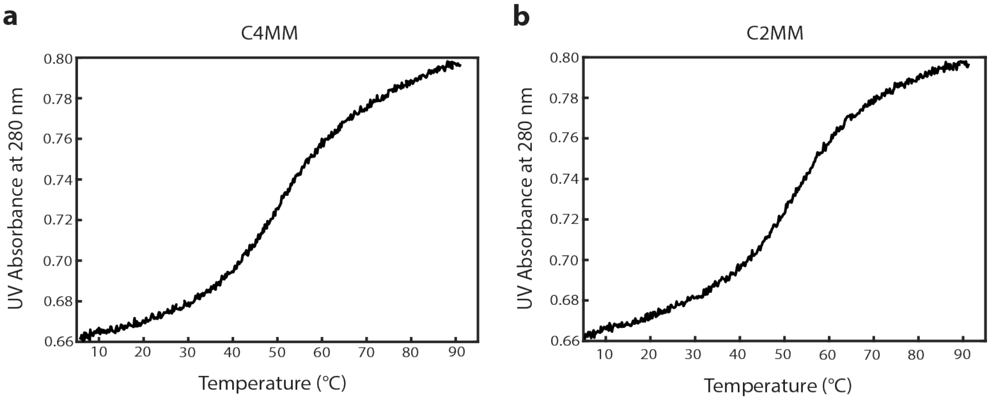


**Figure S3. UV melting experiments for C4MM and C2MM.** UV melting curves for **(a)** C4MM and **(b)** C2MM were collected in 100 mM NaCl, 10 mM $NaH_2PO_4/Na_2HPO_4$, pH = 7.0.

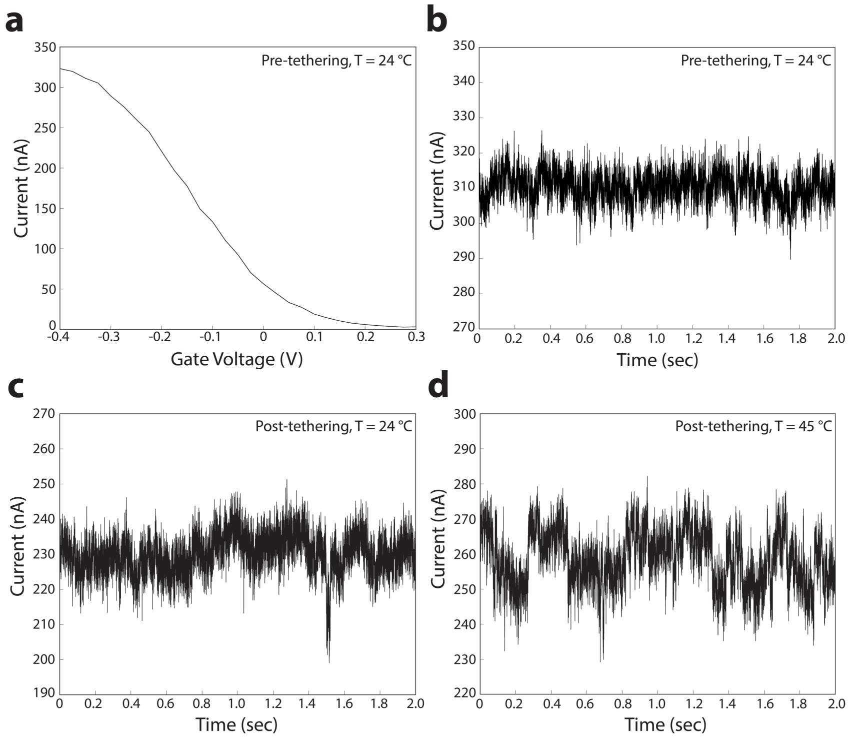


**Figure S4. Temperature dependence of current versus time trajectories recorded using a single, representative smFET device containing a tethered RNA. (a)** I-V curve for this smFET device. Trajectories were recorded: **(b)** at $T_{low}$, prior to the introduction of pyrene-NHS into the flow cell; **(c)** at $T_{low}$, after the incubation of the flow cell with pyrene-NHS and the introduction of a 5'-primary-amine-modified, stem-loop construct; and **(d)** at $T_{high}$ after the recording in (b).

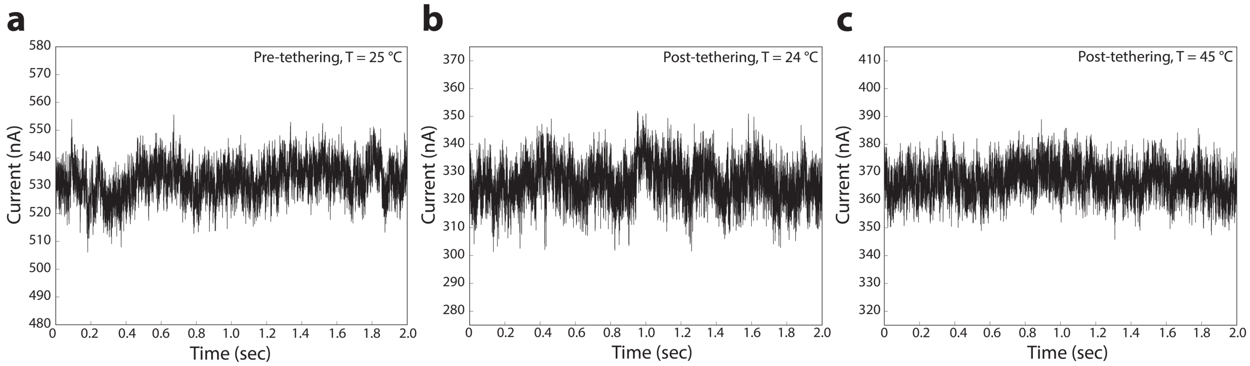


**Figure S5. Temperature dependence of current versus time trajectories recorded using a single, representative smFET device lacking a tethered RNA.** Trajectories were recorded: **(a)** at $T_{low}$, prior to the introduction of pyrene-NHS into the flow cell; **(b)** at $T_{low}$, after the incubation of the flow cell with pyrene-NHS and the introduction of a 5'-primary-amine-modified, stem-loop construct; and **(c)** at $T_{high}$ after the recording in (b).

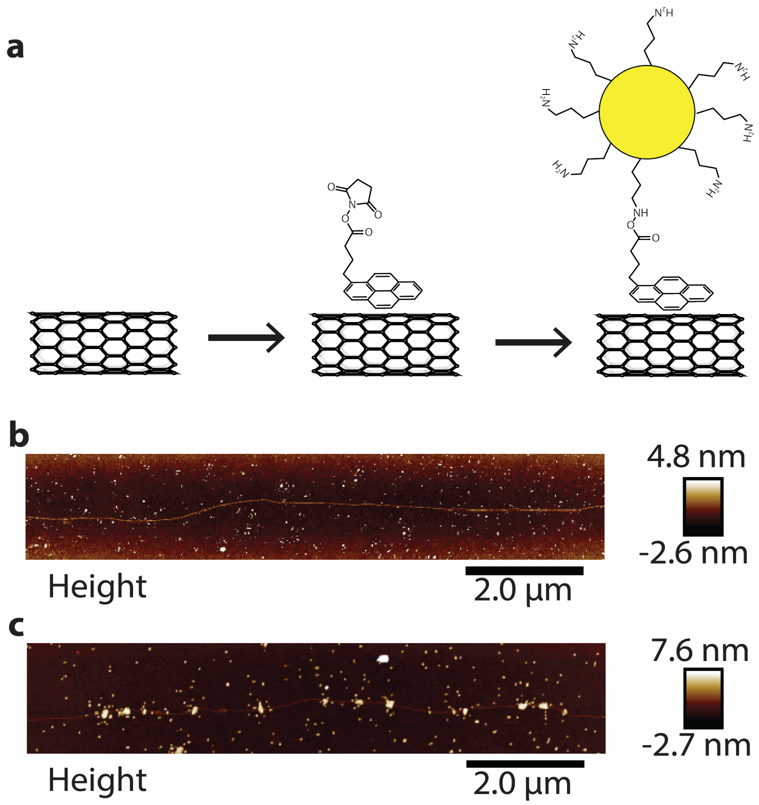


**Figure S6. Validating the specific tethering of pyrene onto SWCNTs. (a)** Schematic cartoon of the non-covalent stacking of the pyrene moiety of pyrene-NHS on the SWCNT surface of an smFET device and the subsequent reaction of the NHS moiety with a primary-amine-functionalized gold nanoparticle. **(b-c)** AFM images of smFET devices and gold nanoparticles after running the reaction depicted in (a) using (b) pyrene-COOH or (c) pyrene-NHS.

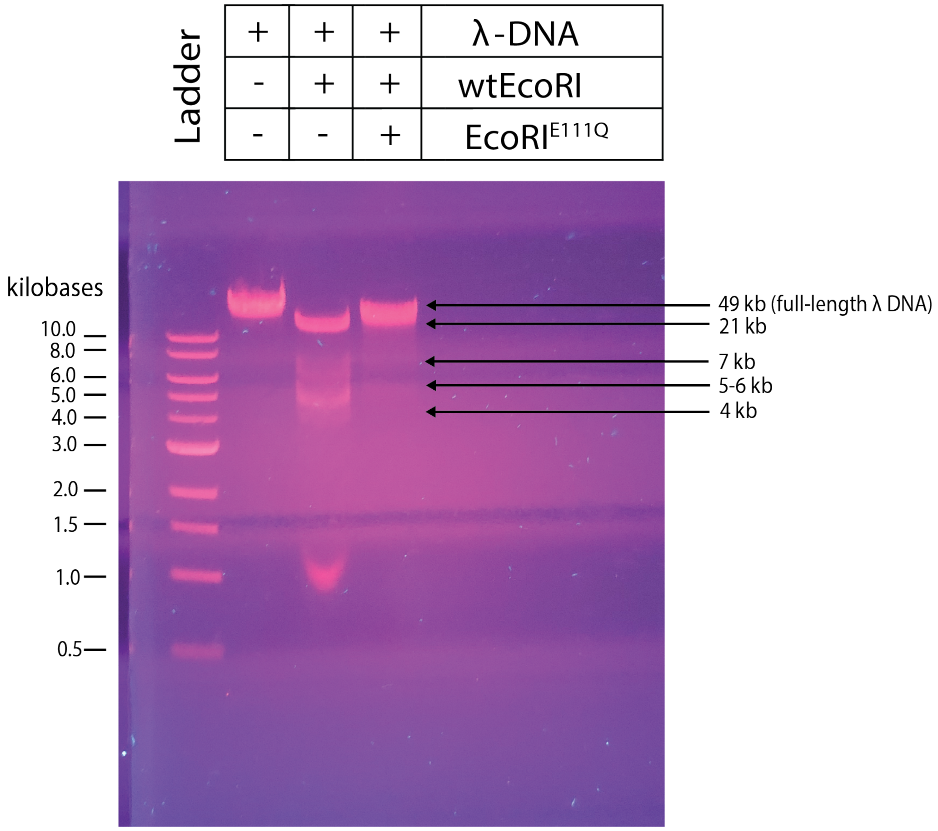


**Figure S7. Characterization of purified EcoRI$^{E111Q}$**. An 0.6 % agarose gel showing the analysis of DNA fragments produced by wtEcoRI-mediated nucleolytic cleavage and protected from wtEcoRI-mediated nucleolytic cleavage by EcoRI$^{E111Q}$ as part of an EcoRI digestion protection assay.

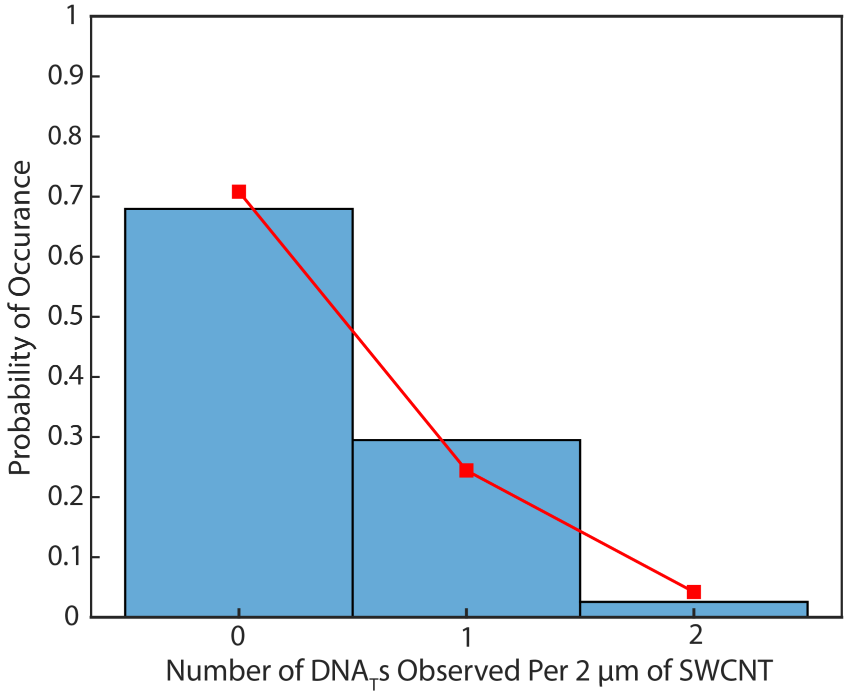


**Figure S8. Distribution of the number of tethered DNA$_T$s per 2µm of SWCNT.** The probabilities of observing 0, 1, or 2 DNA$_T$s, detected using AFM imaging of triple-EcoRI$^{E111Q}$-dimer-bound DNA complexes, tethered per 2µm of SWCNT is plotted as a bar graph. These probabilities are well described by the Poisson distribution function reported in Section I-B and shown as a red datapoints and curve overlaid on the bar graph.

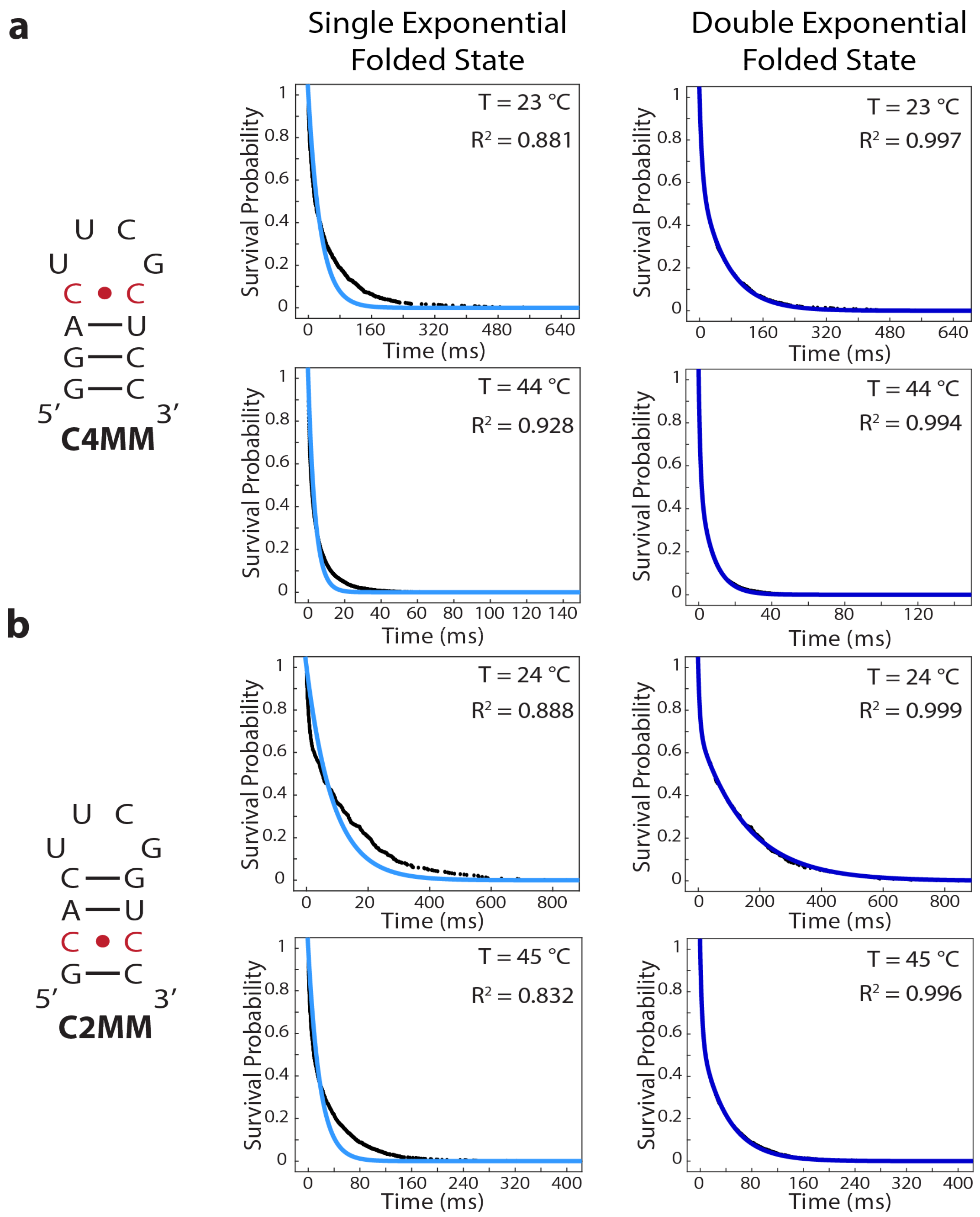


**Figure S9. Single- and double-exponential fitting of survival probability plots of the ensemble of folded conformations.** Secondary structure diagrams of the stem-loop constructs with the mismatched base-pairs denoted in red (first column), single-exponential fits (blue curves) to the survival probability plots of the ensemble of folded conformations (black data points) at $T_{low}$ (second-column, top) and $T_{high}$ (second column, bottom), and double-exponential fits (blue curves) to the survival probability plots of the ensemble of folded conformations (black data points) at $T_{low}$ (third column, top) and $T_{high}$ (third column, bottom) for **(a)** C4MM and **(b)** C2MM.

### III. Supplementary Tables

**Table S1. Thermodynamic parameters for stem-loop formation based on UV melting experiments.**

| Stem-loop Construct | $\Delta H^a$ (kcal mol$^{-1}$) | $\Delta S^a$ (e.u) | $\Delta G_{37}{}^a$ (kcal mol$^{-1}$) | $T_m{}^a$ (°C) |
|---|---|---|---|---|
| C4MM | –29.2(0.7) | –90(2) | –1.22(0.04) | 50.5(0.8) |
| C2MM | –30(2) | –92(5) | –1.47(0.09) | 53.1(0.1) |

[a] Parameters are from denaturation experiments in 100 mM NaCl, 10 mM $NaH_2PO_4/Na_2HPO_4$, pH = 7.0. For UV melting data, parameters are the average of 2 independently prepared samples fit to a two-state model.

**Table S2. RNA sequences of the stem-loop constructs.**

| Stem-loop Construct | Sequence |
|---|---|
| C4MM | 5'-$NH_2(CH_2)_6$ GGA CUU CGC UCC-3' |
| C2MM | 5'-$NH_2(CH_2)_6$ GCA CUU CGG UCC-3' |

**Table S3. DNA sequences used for EcoRI$^{E111Q}$-based DNA tethering controls.**

| DNA Strand | Sequence[a] |
|---|---|
| $DNA_T$ | 5'–$NH_2(CH_2)_6$ GGT GA**G AAT TC**G GCC TGAAAT **GAA TTC** TAA GCG AAG T**GA ATT C**AA ACA–3' |
| $DNA_C$ | 5'–TGT TT**G AAT TC**A CTT CGC TTA **GAA TTC** ATT TCA GGC C**GA ATT C**TC ACC–3' |

[a] EcoRI$^{E111Q}$ binding sites are bolded and underlined

**Table S4. Comparison of thermodynamic parameters determined from smFET and UV melting experiments.**

| Construct | T[a] (°C) | smFET ΔG (kcal mol$^{-1}$) | UV melting ΔG (kcal mol$^{-1}$) | smFET $f_u$ | UV melting $f_u$ |
|---|---|---|---|---|---|
| C4MM | 23 | –1.20(0.06) | –2.48(0.01) | 0.11 | 0.02 |
| | 44 | –0.4(0.2) | –0.59(0.06) | 0.33 | 0.28 |
| C2MM | 24 | –1.8(0.2) | –2.7(0.2) | 0.05 | 0.01 |
| | 45 | –0.9(0.1) | –0.74(0.05) | 0.19 | 0.24 |

[a] ΔG and $f_u$ for smFET experiments were determined at the specified temperatures. Corresponding ΔG and $f_u$ for UV melting experiments were calculated from the standard thermodynamic parameters in Table S1.

**Table S5: Overall folding and unfolding rate constants of UNCG tetraloops derived from previous T-jump studies using a sequential four-state model.**

| Sequence | $T^a$ (°C) | $k_f$ ($s^{-1}$) | $k_u$ ($s^{-1}$) | $\tau_f$ (ms) | $\tau_u$ (ms) |
|---|---|---|---|---|---|
| gacUUCGguc[b] | 55 | 7223 | 43 | 23.11 | 0.14 |
| gacUACGguc[c] | 55 | 1228 | 458 | 2.18 | 0.81 |

[a] Rate constants are calculated at the given temperature.
[b] Energy parameters are from reference 14.
[c] Energy parameters are from reference 15. Rate constants are calculated based on the SI Materials and Methods section G.

**Table S6: Individual rate constants of UNCG tetraloops derived from previous T-jump studies using a sequential four-state model.**

| Sequence | $T^a$ (°C) | $k_{12}$ ($s^{-1}$) | $k_{23}$ ($s^{-1}$) | $k_{23}$ ($s^{-1}$) | $k_{32}$ ($s^{-1}$) | $k_{34}$ ($s^{-1}$) | $k_{43}$ ($s^{-1}$) |
|---|---|---|---|---|---|---|---|
| gacUUCGguc[b] | 55 | 96290 | 90981 | 26674 | 22537 | 115 | 14366 |
| gacUACGguc[c] | 55 | 91167 | 40681 | 18632 | 12748 | 950 | 1642 |

[a] Rate constants are calculated at the given temperature.
[b] Energy parameters are from reference 14.
[c] Energy parameters are from reference 15. Rate constants are calculated based on the SI Materials and Methods section G.
